# Stable excitatory-inhibitory synapse balance despite dynamic turnover

**DOI:** 10.1101/2025.06.02.657384

**Authors:** Krassimira A. Garbett, James P. Allen, Jaybree M. Lopez, Cassandra M. Smith, Richard C. Sando

## Abstract

Diverse synaptic connections self-organize into neural circuits during brain development. A balance between excitatory and inhibitory synaptic function is required for information processing by these neural circuits. Despite the importance of this balance, the interplay between excitatory and inhibitory synaptic assembly during circuit establishment remains unclear due to a lack of means to monitor both processes simultaneously. Here, we develop imaging and analysis methods to visualize and track excitatory and inhibitory synapses. By applying these approaches, we find that despite continual dynamics, excitatory and inhibitory synaptic density remain at steady-state levels during synapse maturation. These results indicate balanced excitatory and inhibitory synapse assembly despite continual synaptic turnover.

## Introduction

Synapse assembly, maturation, and elimination require spatial and temporal coordination of several cell biological processes. These events are initiated by pre- and postsynaptic recognition of combinatorial networks of cell surface receptors, adhesion molecules, and secreted factors^1^. These extracellular networks subsequently initiate bi-directional signaling cascades that stabilize nascent synapses and recruit the corresponding functional machinery essential for diverse physiological synaptic parameters. Synapse elimination likely involves termination of these signals and disassembly of trans-synaptic adhesion complexes^1,2^. While increasing evidence supports this model, the dynamics of mammalian synapse assembly, together with the interplay between excitatory and inhibitory synapses during these processes, remain incompletely understood due to the paucity of approaches to visualize these events in real time^3^.

A dynamic relationship between excitatory and inhibitory synaptic function is critical for computation by neural circuits. Developmental imbalances between excitatory and inhibitory synapses contribute to neurological disorders, including autism^4^. Moreover, the pathophysiology of Rett syndrome, a severe neurodevelopmental disorder, involves a reduction in spontaneous activity caused by an increase in synaptic inhibition over excitation^5^. Interestingly, a structural coordination between excitatory and inhibitory synapses and functional interactions between these synapses can maintain a constant excitatory/inhibitory ratio along the same dendritic tree^6^. This raises the hypothesis that excitatory and inhibitory synapse assembly is coordinated to maintain a functional balance during development.

Testing this hypothesis requires approaches to simultaneously visualize interactions between mammalian pre- and postsynaptic compartments over relatively long timescales with high spatiotemporal resolution^7–9^. Long term imaging studies have revealed dynamic structural changes associated with circuit remodeling during development. Invertebrate systems enable global and long term time-lapse imaging of circuit assembly *in vivo*^10,11^, including high content, long term visualization of growth cones and filipodial dynamics during *Drosophila* visual system development^12,13^. In mammalian systems, dendritic spines, structures associated with excitatory postsynaptic compartments, can also be visualized long-term during their formation and dynamics^14^. Studies in the CA1 region of live mice provided evidence for a remarkably dynamic turnover of dendritic spines^15^. This was in contrast to studies in the neocortex, which found a higher level of spine stability, suggesting that different cell types may display different synaptic turnover rates to support their network functions^16–18^. Despite these advances, methods to study both excitatory and inhibitory synapses concomitantly are limited.

To address the dynamic interplay between excitatory and inhibitory synapse assembly in mammalian neurons, we generated long-term time-lapse imaging approaches and a computational analysis pipeline to simultaneously monitor the dynamics of presynaptic Synaptobrevin-2/VAMP2 (Syb2), excitatory postsynaptic Homer1c, and inhibitory postsynaptic Gephyrin, which are routinely used to label pre- or postsynaptic compartments. These strategies enable examination of the spatial and temporal dynamics of excitatory and inhibitory synaptic compartments, and their co-clustering into contacts. By applying these approaches, we observed that subsets of synapses are forming and eliminating even in mature hippocampal neurons. Despite this, excitatory and inhibitory synapse levels remain balanced over time. Collectively, these approaches provide new insights into the dynamics and balance of inhibitory and excitatory synapses during mammalian circuit establishment.

## Results

While canonical synaptic proteins are routinely used as markers of pre- or excitatory postsynaptic compartments, long-term live imaging of both simultaneously has been challenging. Thus, we developed live imaging procedures to visualize excitatory synapse dynamics over relatively long timescales (**Fig. 1**). We first generated a set of lentiviral reporters based on routinely used synaptic markers, including mClover3-Homer1c and HaloTag-Syb2 fusion constructs. mClover3 is a brighter and more photostable version of the green fluorescent protein Clover^19^, and HaloTag is a self-labeling tag with a panel of versatile and highly stable fluorescent dyes available, including far red dyes^20^. To test if these reporters could be used for labeling synapses, we validated their proper localization via co-staining hippocampal neurons for other endogenous synaptic markers (**Fig. 1A-F**). Hippocampal neurons transduced with HaloTag-Syb2 were live labeled with cell-permeant JF646 HaloTag ligand^21^, and subsequently immunolabeled for endogenous synaptic markers. As expected, HaloTag-Syb2 co-localized with presynaptic Syn1/2 (Synapsin1/2), and co-clustered together with postsynaptic Homer1 (**Fig. 1A-C**). Additionally, mClover3-Homer1c exhibited higher co-localization with the excitatory postsynaptic marker, SHANK2, compared to the inhibitory postsynaptic marker Gephyrin (**Fig. 1D-F**).

**Figure 1:**
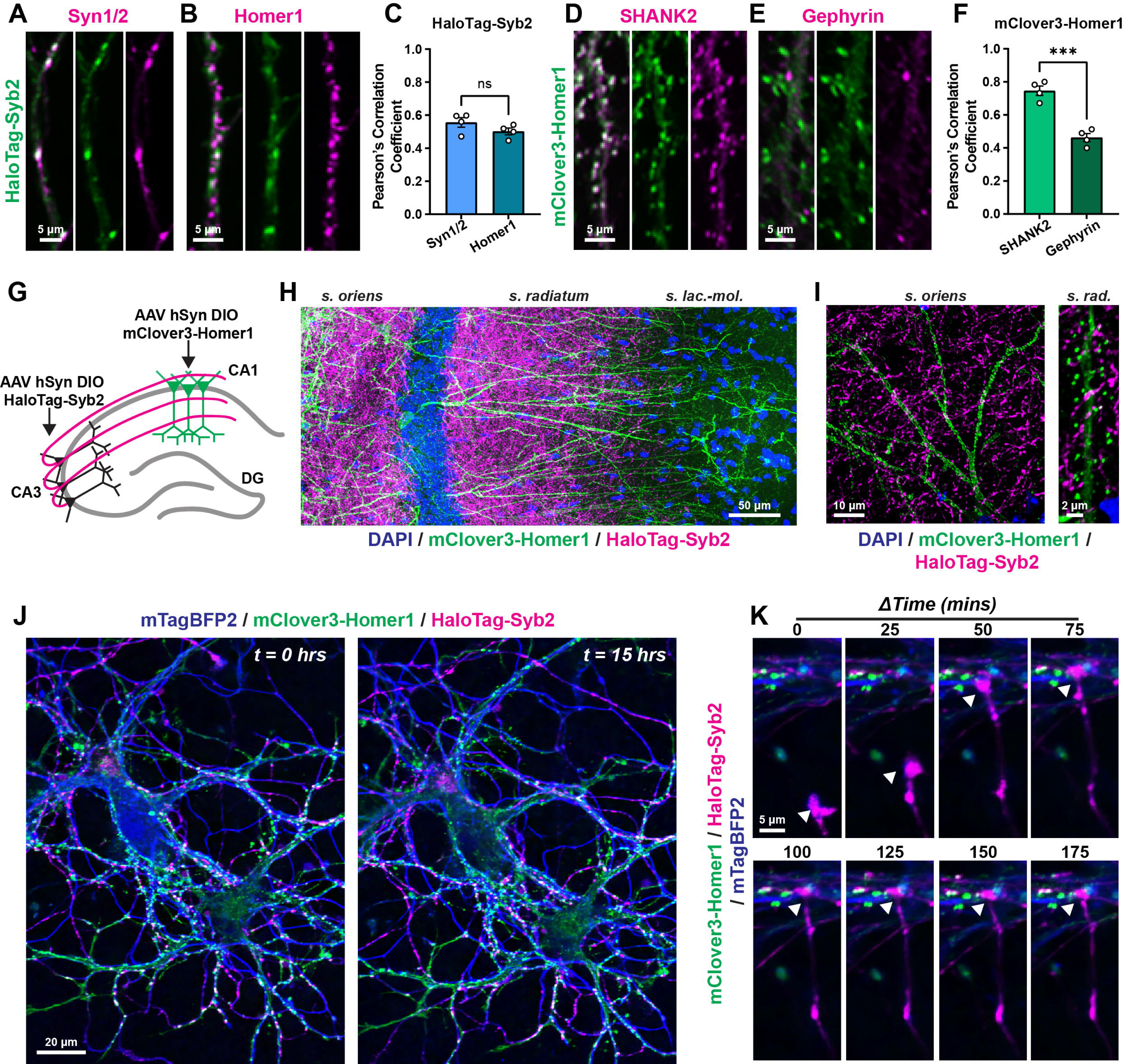
Characterization of live imaging reporters to simultaneously monitor presynaptic and excitatory postsynaptic compartments. **A & B**, lentiviral-delivered HaloTag-Syb2 in primary hippocampal neurons labeled with cell-permeant JF646 HaloTag ligand and subsequently immunostained for Syn1/2 (**A**; presynaptic) or Homer1 (**B**; excitatory postsynaptic). **C,** quantification of Pearson’s correlation coefficient from data in Figure 1A & B. **D & E,** representative dendrites from primary hippocampal neurons transduced with lentivirus encoding mClover3-Homer1c immunostained for SHANK2 (**D**; excitatory postsynaptic) or Gephyrin (**E**; inhibitory postsynaptic). **F,** quantification of Pearson’s correlation coefficient from experiments in Figure 1D & E. **G,** diagram of experimental strategy to label Schaffer collateral synapses *ex vivo*. **H**, representative viral labeling in the CA1 region from experiments outlined in Figure 1G. **I**, high magnification images of the indicated sub-regions of CA1 neurons expressing mClover3-Homer1 and receiving HaloTag-Syb2-positive CA3 Schaffer collateral inputs. **J,** representative live primary hippocampal neuron cultures expressing mClover3-Homer1c, HaloTag-Syb2 and mTagBFP2 before (*left*) and after (*right*) a 15-hr imaging period. **K,** visualization of nascent excitatory synapses in primary cultures. Representative HaloTag-Syb2-labeled growth cone forming a co-cluster with a mClover3-Homer1c punctum along a mTagBFP2-filled dendrite. Numerical data are means ± SEM from 4 independent biological replicates. Statistical significance was assessed with a two-tailed t-test (***, p<0.001). See Movie S1 for representative live imaging.

We next tested their efficacy *ex vivo* in the hippocampal CA1 circuit (**Fig. 1G-I**). We injected AAV encoding HaloTag-Syb2 into the CA3 to label Schaffer collateral inputs into the CA1, and injected AAV expressing mClover3-Homer1c into the CA1. This approach yielded robust labeling of pre- and postsynaptic Schaffer collateral synapses in the stratum oriens and radiatum regions (**Fig. 1G-I**). These results support that our fluorescently-labeled reporters serve as reliable proxies for their respective synaptic compartments in native circuits and demonstrates their utility to label pre- and postsynaptic compartments simultaneously.

We next tested their compatibility and efficacy for long-term time-lapse imaging. Pre- and postsynaptic combinations (HaloTag-Syb2 with mClover3-Homer1c) were lentiviral-transduced into primary hippocampal neurons together with lentiviral-delivered mTagBFP2 as a cell marker. Cells were then labeled live with JF646 HaloTag ligand as before and imaged at DIV12-14, a point when excitatory synapses mature in primary hippocampal neurons^22,23^. Cells were imaged for 15-hrs total at 5-min intervals in a live cell incubation chamber with resonance scanning confocal microscopy (**Fig. 1J & K, Movie S1**). This permitted stable imaging of labeled synaptic compartments, including co-clustered pre- and postsynaptic compartments, without detectable photobleaching (**Fig. 1J**). This imaging strategy captured nascent synapses assembling even in relatively mature primary hippocampal neurons, such as HaloTag-Syb2-positive growth cones approaching and forming contacts with mClover3-Homer1c postsynaptic compartments along mTagBFP2-filled dendrites (**Fig. 1K**).

We next generated custom-built analysis pipelines to enable identification, tracking, and categorization of individual puncta and co-clustered puncta over time (**Fig. 2, Fig. S1, S2**). Due to potential microscope drift over long imaging periods, we created a new drift correction algorithm that corrects for image drift while preserving biological motion (e.g. neurite motion over time) (**Movie S2, see Methods**). We observed that some neurites were highly mobile during the 15-hr imaging period while others were stable. Thus, we binned neurons into two categories, moving or still, based on several quantitative parameters (**Fig. S1**). Puncta from still neurites displayed significantly longer durations, lower speed, and less total net displacement over time compared to moving neurites. (**Fig. S1**). We focused on analysis of still neurites for further tracking and quantification for simplicity.

**Figure 2:**
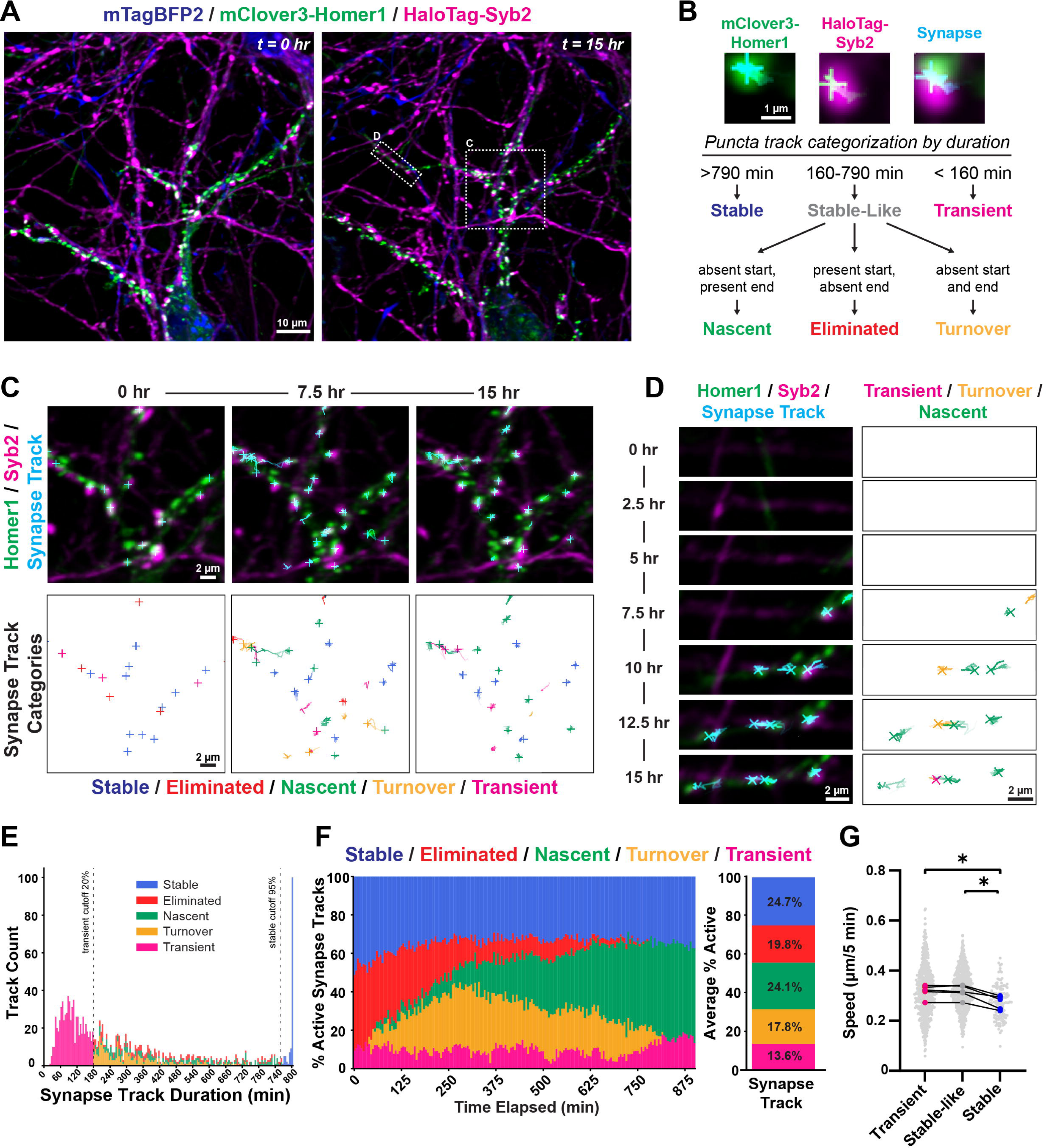
Visualization of distinct populations of excitatory synapses. **A**, live hippocampal cultures expressing mClover3-Homer1c, HaloTag-Syb2, and mTagBFP2 before (*left*) and after (*right*) a 15-hr imaging period. Boxes indicate zoomed-in regions shown in panels C and D. **B,** example tracked pre- and postsynaptic HaloTag-Syb2 and mClover3-Homer1c puncta and synapse tracks. Cyan crosshairs mark current object position, while cyan trails indicate previous 30 frames (5 mins per frame). Tracks were further classified as indicated in the flow chart. **C,** *Top,* representative puncta tracking of co-clustered HaloTag-Syb2 (magenta) and mClover3-Homer1c (green) with synapse tracks (cyan) over 15 hrs. *Bottom,* synapse categories for the same tracks shown. Trails indicate previous 30 frames (2.5 hrs). **D,** as in C, with nascent synapses along an extending dendrite. **E,** histogram of synapse track durations with color-coded track categories. **F**, *Left*, data averaged across time and replicate cultures showing total relative abundance of each track type. *Right,* percentage of each synapse track category at each frame during the imaging session. Each bar represents synapses active at each sampling interval (5 min). N=5 independent cultures. **G,** speed of track motion for Transient, Stable-like, and Stable synapses. One-way RM ANOVA (P=0.0031) with Tukey’s post hoc. N=5 independent cultures. Gray dots are individual track speed; colored points are average track speed per culture. (*, P < 0.05). Post hoc test comparisons were made between all groups, but only comparisons where adj P < 0.05 are shown. See Movie S2 and S3 for representative tracking examples and drift correction. See Figures S1 and S2 for additional characterization of tracking approaches.

To better understand the temporal dynamics of synaptic puncta in cultures, we further stratified synaptic puncta into subclasses. Co-clustered, paired HaloTag-Syb2/mClover3-Homer1c puncta (defined as synapses) were identified and categorized based on their persistence and formation/elimination during the imaging period (**Fig. 2A & B**). Stable synapses persisted over the entire imaging period, while Transient synapses were short-lived. A third intermediate class of synapses were composed of either Nascent, Eliminated, or synapses that Turnover based on their presence and/or absence at the beginning and/or end of imaging (**Fig. 2B**). These analysis tools enabled long-term tracking and categorization of synapses over a 15-hr period (**Fig. 2C & D, Movie S3**). Moreover, they allowed quantification of the abundance of distinct synapse populations and their distribution during the imaging period (**Fig. 2E & F**). Accordingly, we observed sub-populations of synapses being eliminated over time, while others were newly formed (nascent) using these approaches (**Fig. 2F**). Stable synapses displayed significantly lower movement speed compared to transient or stable-like synapses from the same cultures (**Fig. 2G**), suggesting a relationship between temporal and spatial dynamics. These methods were also effective at tracking and quantifying unpaired HaloTag-Syb2 or mClover3-Homer1c puncta (**Fig. S2**). These computational approaches highlight the dynamic nature of excitatory synapses over these timescales and the interplay between pre- and postsynaptic compartments in live hippocampal neurons.

Previous studies have shown that excitatory synapses are functionally maturing at approximately DIV12 in primary hippocampal neurons^22,23^. Therefore, we next examined excitatory synapses relatively early (DIV8-9) and later (DIV11-14) during maturation in culture (**Fig. 3, Fig. S2F & G**). Temporal dynamics of synapses were similar between groups. (**Fig. 3A-F**). Homer1c puncta displayed significantly higher total durations at DIV8-9 compared to presynaptic Syb2 or synapses, suggesting postsynaptic compartments are less mobile and less dynamic relative to presynaptic (**Fig. 3C, S2B, C**). Similarly, at DIV11-14, Homer1 puncta were more persistent than synapse pairs (**Fig. 3D-F**). However, the duration of Syb2 puncta was not significantly different than Homer1c in this group, suggesting that presynaptic compartments become more stable over time (**Fig. 3D-F**).

**Figure 3:**
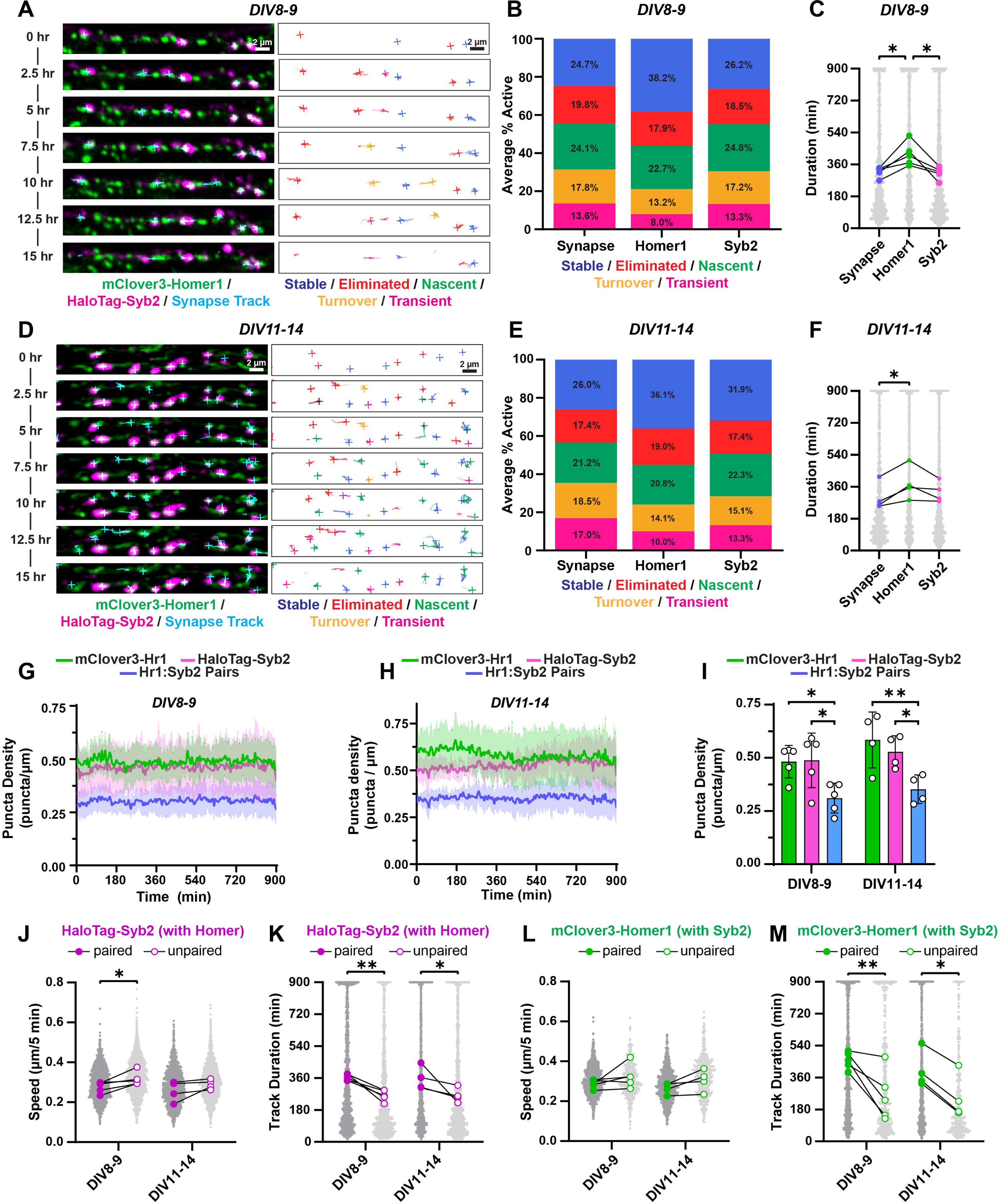
Spatiotemporal dynamics of excitatory synapse populations during maturation. **A**, representative dendrites at DIV8-9 with HaloTag-Syb2 and mClover3-Homer1 with overlayed tracks (*left*) and track categorizations (*right*). Track trails indicate previous 30 frames (2.5 hrs). **B,** breakdown of synapse category averages per frame at DIV8-9 for synapse tracks, mClover3-Homer1c tracks, and HaloTag-Syb2 tracks. Synapse tracks are reproduced from Figure 2F for comparison. **C,** duration of synapse tracks at DIV8-9, mClover3-Homer1c tracks, and HaloTag-Syb2 tracks. One-way RM ANOVA (P=0.001) with Tukey’s post hoc. N=5 independent cultures. Gray dots are individual track duration; colored points are average track duration per culture. **D,** as in A, but with DIV11-14 cultures. **E**, as in B, but with DIV11-14 cultures. **F**, as in C, but with DIV11-14 cultures. One-way RM ANOVA (P=0.03) with Tukey’s post hoc. N=4 independent cultures. **G,** puncta density of mClover3-Homer1c tracks, HaloTag-Syb2, and paired puncta (synapses) across the imaging period for DIV8-9 cultures. Data are Mean ± SD. **H**, as in G, but with DIV11-14 cultures. **I**, average puncta density of DIV8-9 and DIV11-14 cultures. Two-way ANOVA (DIV, P=0.114; Puncta type, P=0.0004; interaction, P=0.74) with Tukey post hoc. **J,** puncta speed of paired and unpaired tracks for HaloTag-Syb2 paired with mClover3-Homer1c. Two-way RM ANOVA (DIV, P=0.31; Pairing, P=0.0116; interaction, P=0.73) with Fisher’s LSD post hoc. N=4-5 independent cultures. Gray dots are individual tracks speed and colored points are average track speed per culture. **K**, track duration of paired and unpaired tracks for HaloTag-Syb2 paired with mClover3-Homer1c. Two-way RM ANOVA (DIV, P=0.78; Pairing, P=0.029; interaction, P=0.82) with Fisher’s LSD post hoc. N=4-5 independent cultures. Gray dots are individual tracks duration and colored points are average track duration per culture. **L,** as in J, but for mClover3-Homer1c paired with Syb2. Two-way RM ANOVA (DIV, P=0.41; Pairing, P=0.0462; interaction, P=0.94) with Fisher’s LSD post hoc. N=4-5 independent cultures. **M,** as in K, but for mClover3-Homer1c paired with Syb2. Two-way RM ANOVA (DIV, P=0.63; Pairing, P=0.0006; interaction, P=0.49) with Fisher’s LSD post-hoc. N=4-5 independent cultures. Post hoc test comparisons were made between all groups, but only comparisons where adj P < 0.05 are shown for brevity. (*, P<0.05, ** P<0.01).

Interestingly, despite these spatiotemporal dynamics, total puncta density remained constant over the 15-hr imaging session in both DIV8-9 and DIV11-14 neurons (**Fig. 3G & H**). Furthermore, the density of synapse pairs was significantly lower than individual pre- and postsynaptic components at both time points, suggesting a subset of available pre- and postsynaptic sites remain even at the later maturity stage (**Fig. 3I**). Many of these available pre-or postsynaptic compartments remain unassociated for the observable period, while other puncta tightly co-associated for a prolonged time (**Fig. S2D**). Based on that we categorized them as either unpaired or paired (**Fig. S2E**). We then quantified the speed and track duration of paired and unpaired puncta at the two different time points (**Fig. 3J-M**). Unpaired Syb2 puncta displayed higher speed than paired in DIV8-9 cultures compared to DIV11-14, supporting more highly dynamic presynaptic compartments are present early during maturation (**Fig. 3J**). Unpaired Syb2 track durations were significantly shorter than paired in both culture periods, suggesting paired puncta persist longer (**Fig. 3K**). However, unpaired Homer1c puncta speed was comparable to paired, indicating less mobility relative to presynaptic sites (**Fig. 3L**).

Paired Homer1c puncta again persisted longer than unpaired in both culture periods (**Fig. 3M**). We also observed similar levels of colocalization and similar total amounts of puncta in both culture periods (**Fig. S2F-G**). These results highlight that neurons maintain an equilibrium of excitatory synapse density despite continual spatiotemporal dynamics.

We then generated analogous lentiviral tools to facilitate visualization and tracking of inhibitory synapses over time (**Fig. 4**). To facilitate compatibility with our other tools, we generated a tdTomato-Gephyrin fusion protein. We validated that this fluorescent fusion protein was highly co-localized with the inhibitory postsynaptic marker GABARα1 but not the excitatory postsynaptic marker Homer1 (**Fig. 4A**). We then tested the efficacy of the AAV-delivered fusion protein in the hippocampal CA1 region (**Fig. 4B & C**). We utilized the Chrna2-Cre line, which is selective for hippocampal OLM (oriens lacunosum-moleculare) interneurons^24,25^. These neurons reside in the stratum oriens and project to the stratum lacunosum-moleculare (**Fig. 4B**). Contrary to our experiments labeling excitatory synapses in the stratum oriens and radiatum (**Fig. 1G-I**), this approach selectively labeled inhibitory synapses in the stratum lacunosum-moleculare (**Fig. 4B & C**). These results support the fidelity of these tools in labeling specific excitatory or inhibitory postsynaptic compartments.

**Figure 4:**
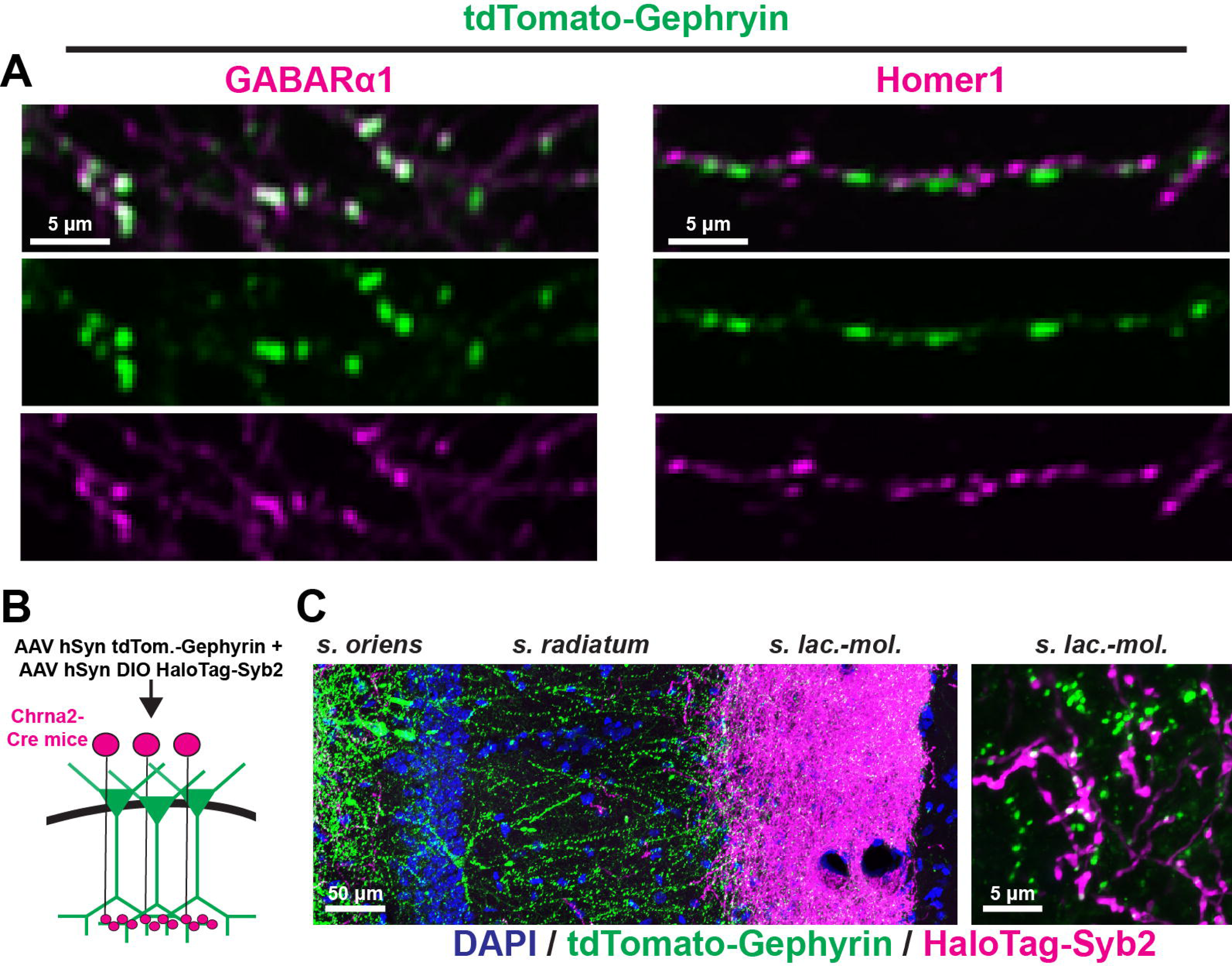
Labeling approaches for inhibitory synapses *in vitro* and *ex vivo*. **A**, co-localization of the tdTomato-Gephyrin lentiviral reporter with GABARα1 (inhibitory postsynaptic; *left*) or Homer1 (excitatory postsynaptic; *right*). **B,** diagram of experimental strategy to virally label OLM interneuron-CA1 synapses. **C,** representative CA1 neurons expressing tdTomato-Gephyrin and receiving HaloTag-Syb2-positive presynaptic terminals from OLM interneurons. *Right*, high magnification image of the stratum lacunosum-moleculare region.

We then co-transduced primary hippocampal neurons with tdTomato-Gephyrin, HaloTag-Syb2, and mTagBFP2, and conducted live imaging, tracking, and quantifications as before, again comparing DIV8-9 to DIV11-14 cultures (**Fig. 5, S3, Movie S4**). Our analysis tools effectively visualized presynaptic and inhibitory postsynaptic puncta, as well as co-clustered pairs, over the 15-hr imaging period, and captured dynamic populations of inhibitory synapses (**Fig. 5A-F**). Postsynaptic Gephyrin puncta trended towards more stable at both culture maturation times, with a larger fraction of stable synapse subtypes (**Fig. 5B & E**), but no significant difference in duration from Syb2 (**Fig. 5C & F**). Additionally, synapse tracks of co-clustered puncta pairs were shorter in duration than individual Syb2 tracks (**Fig. 5C & F**), suggesting inhibitory postsynaptic compartments may not associate as strongly with Syb2-containing pre-synapses as excitatory postsynaptic compartments. This was consistent with other measurements including broader distribution of inter-puncta distances (**Fig. S2D, Fig. S3D)**, larger fraction of unpaired Syb2 (**Fig. S2E, Fig. S3E)**, and low fraction of paired puncta and co-cluster density (**Fig. S2F, G, Fig. S3F, G**). Furthermore, we saw continuous turnover of puncta that maintained a stable density along dendrites (**Fig. 5G-I**). We saw no significant differences between inhibitory paired and unpaired puncta motion speed (**Fig. 5J & L**). However, Gephyrin synaptic pairs were substantially longer in duration when paired with Syb2 compared to non-paired Gephyrin puncta (**Fig. 5M**), but not Syb2 paired with Gephyrin (**Fig. 5K**). This observation suggests the Gephyrin postsynaptic association with Syb2 pre-synapses drives its puncta stability, but Syb2 puncta stability is likely primarily driven by association with excitatory postsynaptic compartments as observed previously (**Fig. 3J & K**). Altogether, these tools enable long-term visualization and quantitative assessment of inhibitory synapses in primary hippocampal neurons.

**Figure 5:**
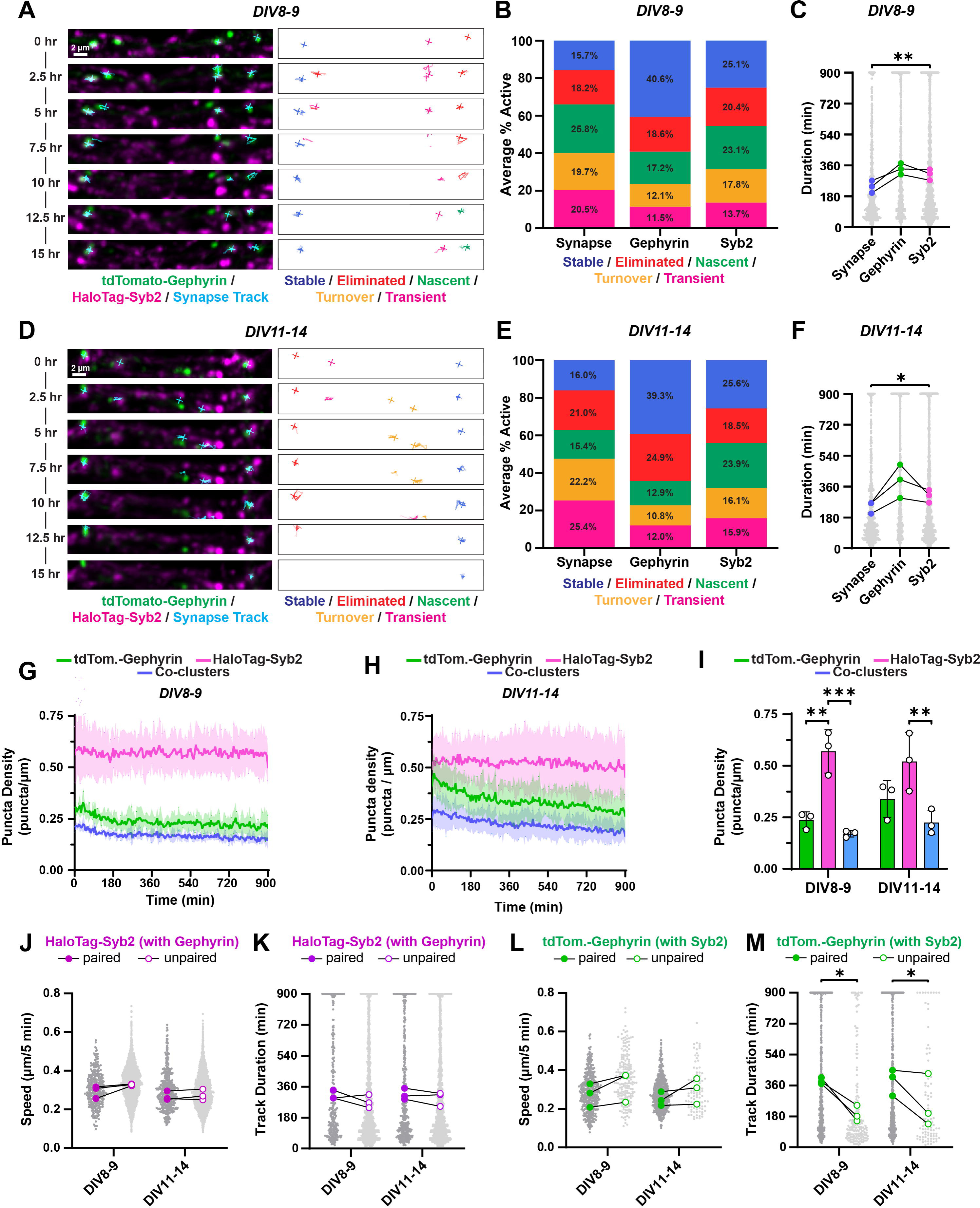
Live imaging inhibitory synapse spatiotemporal dynamics. **A**, representative dendrites at DIV8-9 with HaloTag-Syb2 and tdTomato-Gephyrin with overlayed tracks (*left*) and track categorizations (*right*). Track trails indicate previous 30 frames (2.5 hrs). **B,** breakdown of synapse category averages per frame across all cultures DIV8-9 for synapse tracks, tdTomato-Gephyrin tracks, and HaloTag-Syb2 tracks. **C,** duration of synapse tracks at DIV8-9, tdTomato-Gephyrin tracks, and HaloTag-Syb2 tracks. One-way RM ANOVA (P=0.0379) with Tukey’s post hoc. N=3 independent cultures. Gray dots are individual tracks duration and colored points are average track duration per culture. **D,** as in A, but with DIV11-14 cultures. **E**, as in B, but with DIV11-14 cultures. **F**, as in C, but with DIV11-14 cultures. One-way RM ANOVA (P=0.09) with Tukey’s post hoc test. **G,** puncta density of tdTomato-Gephyrin tracks, HaloTag-Syb2, and paired puncta (synapses) across the imaging period for DIV8-9 cultures. Data are Mean ± SD. **H**, as in G, but with DIV11-14 cultures. **I**, average puncta density of DIV8-9 and DIV11-14 cultures with tdTomato-Gephyrin and HaloTag-Syb2. Two-way ANOVA (DIV, P=0.39; Puncta type, P<0.0001; interaction, P=0.32) with Tukey post hoc. **J,** puncta speed of paired and unpaired tracks for HaloTag-Syb2 paired with Gephyrin. Two-way RM ANOVA (DIV, P=0.095; Puncta type, P=0.088; interactions P=0.21). **K**, track duration of paired and unpaired tracks for HaloTag-Syb2 paired with Gephyrin. Two-way RM ANOVA (DIV, P=0.63; Puncta type, P=0.15; interaction, P=0.67). **L,** as in J, but for tdTomato-Gephyrin paired with Syb2. Two-way RM ANOVA (DIV, P=0.61, Puncta type, P=0.0592, interaction, P=0.86). **M,** as in K, but for tdTomato-Gephyrin paired with Syb2. Two-way RM ANOVA (DIV, P=0.67; Puncta type, P=0.0084; interaction, P=0.42) with Fisher’s LSD post hoc. Post hoc test comparisons were made between all groups, but only comparisons where adj P < 0.05 are shown. (*, P<0.05, ** P<0.01, *** P<0.001). See Movie S4 for representative live imaging data and Figure S3 for additional characterization of tracking approaches.

Given that our fluorescent reporters rely on overexpression of the tagged proteins, as a parallel approach, we also applied recent advances in CRISPR/Cas9 knock-in strategies to introduce tags into genomic loci within postmitotic neurons^26^ (**Fig. 6, Figs. S4-6**). We employed the TKIT (tandem knock-in with two guides) approach that enables introduction of tags into endogenous neuronal genomic loci with high efficacy^26^. Moreover, the TKIT labeling approach avoids the concern of variable indels in coding regions following CRISPR modification as observed with other strategies^27,28^. Consistent with these studies, we found that this system robustly tagged endogenous GluA2 in primary hippocampal neurons (**Fig. S4A & B**). Since we are focused on developing tagging strategies for commonly used pre- and postsynaptic markers, we developed TKIT constructs for presynaptic Bassoon, excitatory postsynaptic Homer1c, and inhibitory postsynaptic Gephyrin, and generated AAVs encoding the respective sgRNAs/donor and Cas9.

**Figure 6:**
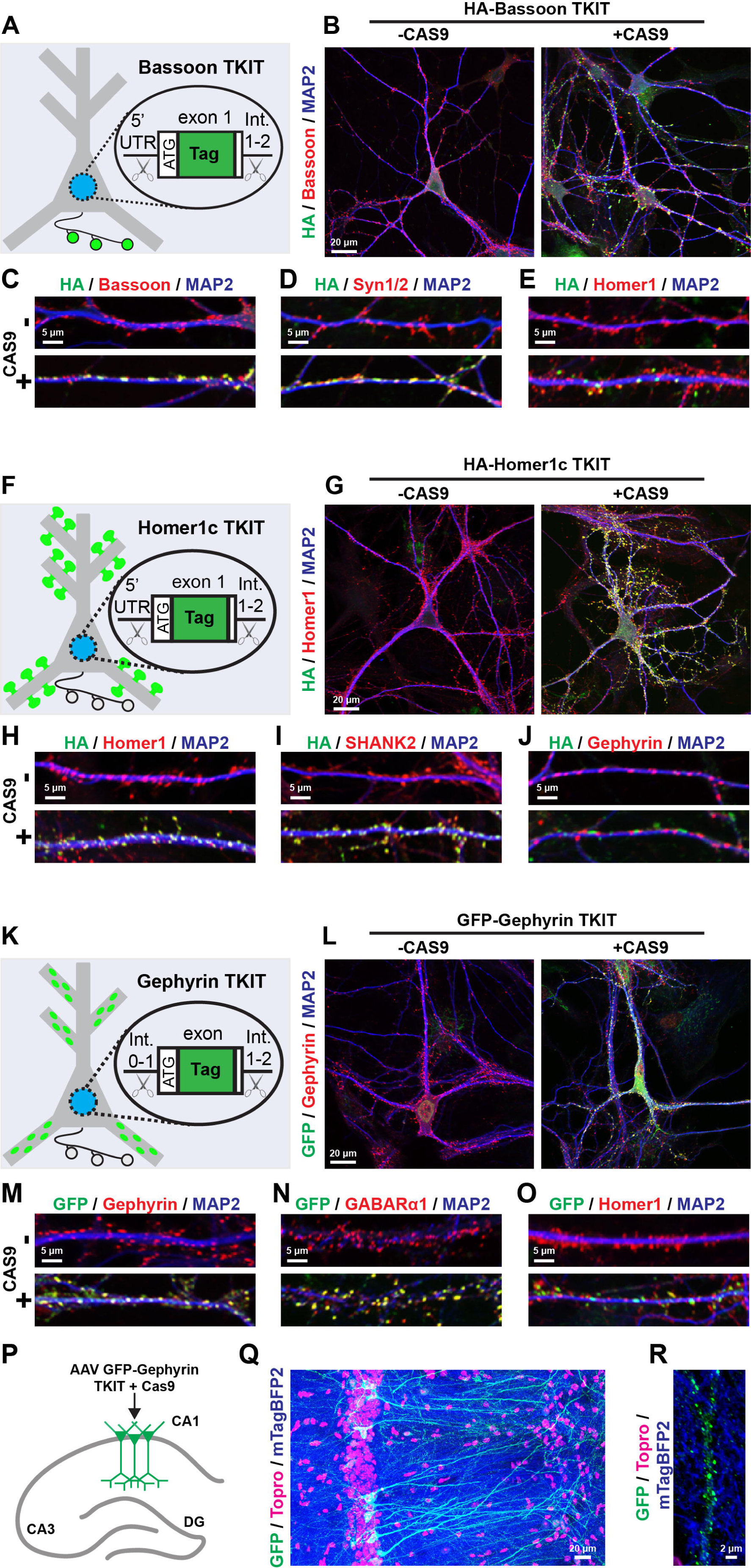
Characterization of TKIT CRISPR/Cas9-based reporters to label endogenous pre- or postsynaptic compartments. **A-E**, characterization of CRISPR/Cas9 tagging strategy for endogenous presynaptic Bassoon. **A,** diagram of TKIT CRISPR/Cas9 approach to tag endogenous Bassoon via AAV-mediated delivery of TKIT CRISPR components. **B,** example neurons transduced with AAVs encoding an HA-Bassoon DNA donor, sgRNAs, without (*left*) or with (*right*) Cas9. Neurons were co-stained for HA, endogenous Bassoon, and the somatodendritic marker MAP2. **C-E,** co-localization of HA-tagged Bassoon with endogenous Bassoon (**C**), another presynaptic marker Syn1/2 (**D**), and the excitatory postsynaptic marker Homer1 (**E**). **F-J,** TKIT tagging of endogenous postsynaptic excitatory Homer1c. **F,** diagram of the CRISPR/Cas9-mediated tagging strategy for endogenous Homer1c. **G,** example primary hippocampal neurons transduced with AAVs encoding the HA-Homer1c DNA donor and sgRNAs without (*left*) or with (*right*) Cas9. Neurons were co-stained for HA tag, endogenous Homer1, and MAP2. **H-J,** co-localization of HA-tagged Homer1c with endogenous Homer1 (**H**), another excitatory postsynaptic marker SHANK2 (**I**), and the inhibitory postsynaptic marker Gephyrin (**J**). **K-O,** validation of TKIT-based tagging of endogenous inhibitory postsynaptic Gephyrin. **K,** diagram of endogenous Gephyrin tagging via TKIT CRISPR/Cas9. **L,** example primary hippocampal neurons transduced with AAVs encoding the Gephyrin tagging donor and sgRNAs without (*left*) or with (*right*) Cas9. Neurons were co-stained for GFP, endogenous Gephyrin, and MAP2. **M-O,** immunostaining for GFP-tagged Gephyrin together with endogenous Gephyrin (**M**), another inhibitory postsynaptic component GABARα1 (**N**), and the excitatory postsynaptic marker Homer1 (**O**). **P-R,** TKIT-mediated Gephyrin tagging in the CA1 region. **P,** diagram of experimental strategy to label endogenous Gephyrin in CA1 neurons. **Q,** GFP tagging of endogenous Gephyrin in the CA1 region. mTagBFP2 was used as an injection site marker. **R,** representative CA1 neuron dendrite labeled with GFP-tagged endogenous Gephyrin. See Figures S4-S6 for additional characterization of TKIT CRISPR/Cas9 tagging approaches.

We first tagged Bassoon in primary hippocampal cultures with an N-terminal HA tag (**Fig. 6A-E, Fig. S5A**). When we virally transduced cultures with HA-Bassoon donor and sgRNAs, we detected robust HA labeling only when Cas9 was also co-delivered. The HA staining overlapped with the staining for endogenous Bassoon at DIV12-14 (**Fig. 6B & C**). Endogenously tagged HA-Bassoon also co-localized with presynaptic Syn1/2, and formed co-clusters with postsynaptic Homer1 (**Fig. 6D, E, Fig. S5A**). Primary neurons harboring TKIT AAVs for HA tagging of Homer1c displayed HA staining that co-localized with endogenous Homer1 only when Cas9 was co-delivered (**Fig. 6G & H**). Furthermore, HA-Homer1c co-localized with another excitatory postsynaptic protein, SHANK2, but was excluded from inhibitory postsynaptic sites stained for Gephyrin (**Fig. 6I, J, Fig. S5B**). We used an analogous approach to tag endogenous Gephyrin with GFP, which was based on the design of Fang *et al*., 2020 but with further optimized sgRNAs. As expected, tagged Gephyrin co-localized to inhibitory synapses with high fidelity and was excluded from excitatory synapses (**Fig. 6K-O, Fig. S5C**). Namely, GFP-tagged Gephyrin co-localized with endogenous Gephyrin and GABARα1, but not excitatory Homer1 (**Fig. 6M-O, Fig. S5C**). The Gephyrin TKIT tagging approach was also effective in the hippocampal CA1 region, where injection of the Gephyrin TKIT components resulted in GFP-Gephyrin fusion throughout CA1 pyramidal neurons (**Fig. 5P-R**). Finally, we found that these tools could label both Homer1c and Gephyrin in the same neuron, indicating the potential to modify two targets in the same cell albeit at low efficacy (**Fig. S5D**).

Tagged mRNA transcripts were only detected when donor/sgRNA AAVs were co-delivered with Cas9 (**Fig. S5E-G**). Moreover, we estimated the overall efficiency of Homer1c and Gephyrin TKIT using standard curve RT-qPCR, and found they modify approximately 5% of total transcripts (**Fig. S5H**). We subsequently assessed potential CRISPR/Cas9 off-target effects (**Fig. S6**). We identified the top two potential off-target sites for each gRNA, PCR amplified the genomic loci and sequenced to detect modifications (**Fig. S6**). For the Homer1c TKIT, sequencing the targeted region detected introduction of the HA tag in the desired location, confirming TKIT modification (**Fig. S6A**). The sequence of the predicted top two off-target sites for each Homer1c sgRNA matched the consensus genomic sequence, supporting the absence of off-target modifications (**Fig. S6B & C**). Similarly, we identified TKIT-mediated tag introduction in Gephyrin and Bassoon without evidence of off-target modifications (**Fig. S6D-I**). These CRISPR/Cas9 approaches build upon the original TKIT toolbox and facilitate the fusion of tags with endogenous presynaptic and postsynaptic proteins.

We subsequently used a Gephyrin donor to label this inhibitory postsynaptic protein with tdTomato for live imaging (**Fig. 7, Fig. S7**). We then conducted time-lapse imaging as described before in CRISPR-modified neuronal cultures at DIV11-14 (**Fig. 7A, Movie S5**). To concurrently monitor presynaptic terminals and endogenous postsynaptic inhibitory compartments, we introduced tdTomato-Gephyrin TKIT AAVs together with lentiviral HaloTag-Syb2 and mTagBFP2 (**Fig. 7B-H, Fig. S7**). As a result, we detected and quantified the distinct subpopulations of synapses (**Fig. 7B-D**). Furthermore, we quantified the individual and the paired HaloTag-Syb2:tdTomato-Gephyrin density along mTagBFP2-labeled dendrites during the imaging period (**Fig. 7E & F)**. As with lentiviral pre- and postsynaptic tracking, a steady-state density of puncta was maintained over time despite dynamic sub-populations (**Fig. 7B-F**). Our analyses tools allowed us to characterize the temporal and spatial properties of inhibitory synapses and the non-paired synaptic compartments in neurons with fluorescently-fused endogenous proteins. As a result, we demonstrated their similarity to the properties observed in cultures with overexpressed reporters (**Fig. 7J-M, S7**). Collectively, this molecular toolbox and tracking approaches enable quantification of the spatiotemporal dynamics and densities of excitatory and inhibitory synapses in live cells.

**Figure 7:**
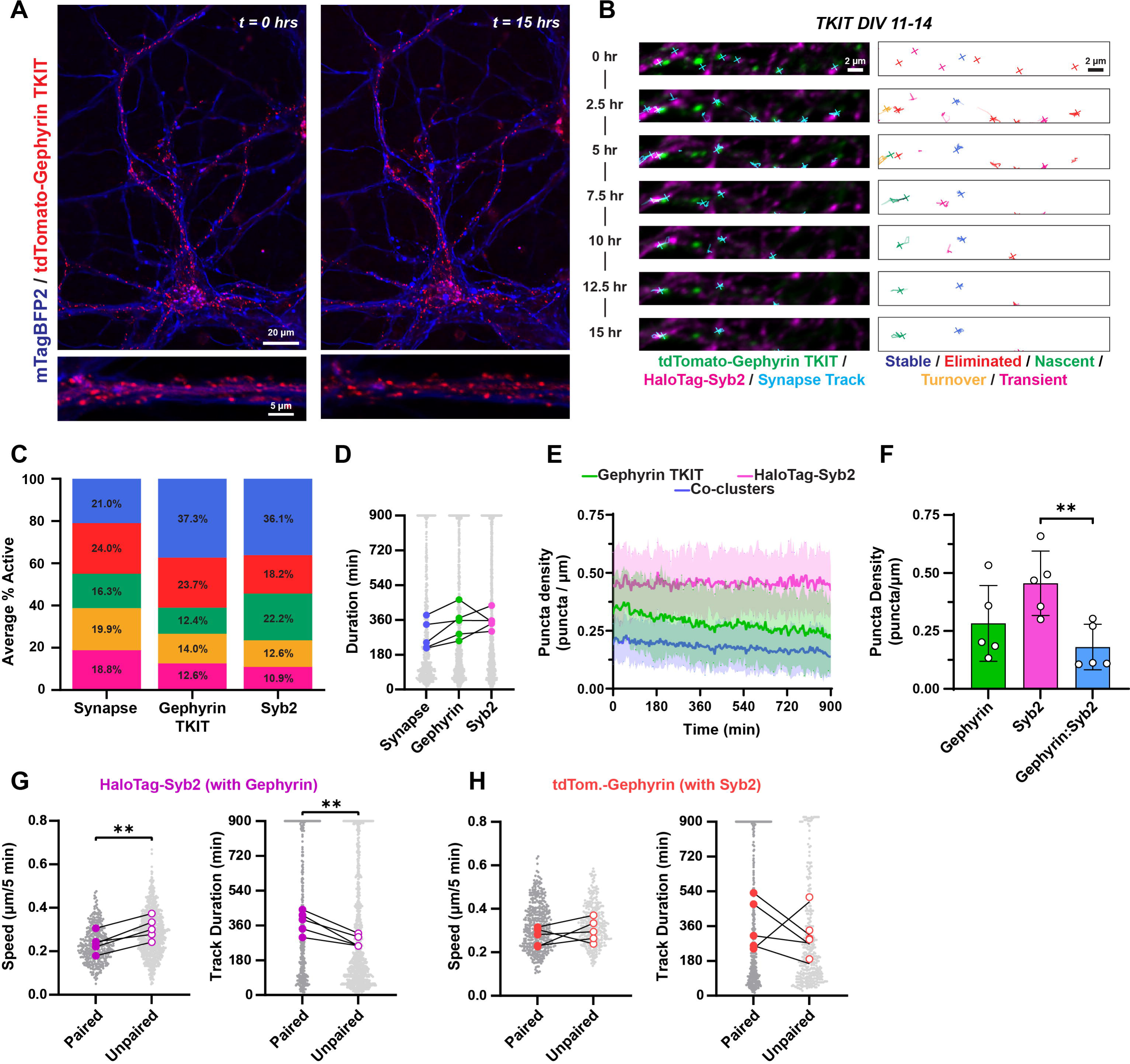
Long-term imaging of endogenous inhibitory postsynaptic Gephyrin. **A**, example primary hippocampal neuron harboring tdTomato-tagged endogenous Gephyrin before and after a 15-hr imaging period. **B,** example dendrite with tdTomato-labeled endogenous Gephyrin and lentiviral expressed HaloTag-Syb2 time-lapse imaged over a 15-hr period, with overlayed tracks (*left*) and track categorizations (*right*). **C,** breakdown of synapse category averages per frame across all cultures DIV8-9 for synapse tracks, tdTomato-Gephyrin tracks, and HaloTag-Syb2 tracks. **D,** duration of synapse tracks, tdTomato-Gephyrin tracks, and HaloTag-Syb2 tracks. One-way RM ANOVA (P=0.1213). N=5 independent cultures. Gray dots are individual tracks duration and colored points are average track duration per culture. **E,** puncta density of endogenous tdTomato-Gephyrin tracks, HaloTag-Syb2, and paired puncta (synapses) across the imaging period. Data are Mean ± SD. **F**, average puncta density of cultures with endogenously labeled tdTomato-Gephyrin and lentiviral HaloTag-Syb2. One-way ANOVA (P=0.01) with Tukey’s post hoc. **G,** puncta speed (*left*) and track duration (*right*) of paired and unpaired tracks for HaloTag-Syb2 paired with endogenous Gephyrin. Paired t-tests, N=5 independent cultures. **H**, as in G, with endogenous tdTomato-Gephyrin paired with HaloTag-Syb2. Paired t-tests, N=5 independent cultures. Post hoc test comparisons were made between all groups, but only comparisons where adj P < 0.05 are shown. (**, P<0.01). See Movie S5 for representative live imaging data and Figure S7 for additional characterization of tracking approach.

We next analyzed excitatory and inhibitory synapses concurrently in neurons transduced with mClover3-Homer1c, tdTomato-Gephyrin, HaloTag-Syb2, and mTagBFP2 lentiviral reporters (**Fig. 8A**). We quantified the density of Homer1c:Syb2 excitatory and Gephyrin:Syb2 inhibitory pairs at DIV8-9 and DIV11-14 (**Fig. 8B & C**). Excitatory synapse density was significantly higher than inhibitory at both times (**Fig. 8B**). We next quantified the ratio of total Homer1c:Syb2 co-clustered puncta counts relative to Gephyrin:Syb2 co-clusters from the same cultures and found that the overall ratio of excitatory to inhibitory synapses remained unchanged over this period (**Fig. 8C**). These results suggest that despite the dynamic environment, hippocampal neurons maintain a balanced equilibrium of excitatory and inhibitory synapses during this period of culture maturity.

**Figure 8:**
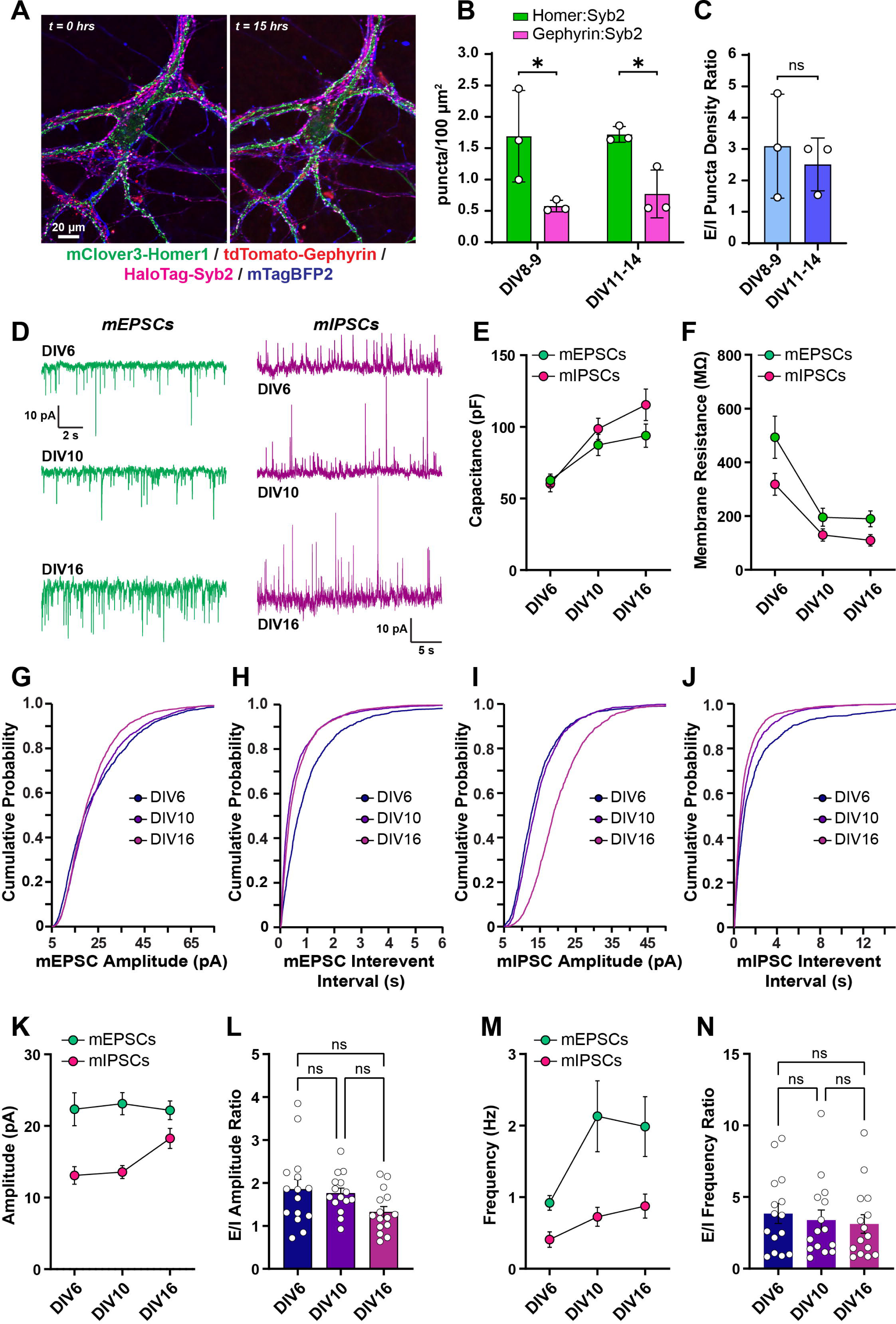
Monitoring excitatory and inhibitory synapse ratios and functional maturation during synaptogenesis. **A**, representative primary hippocampal neurons expressing lentiviral-delivered mClover3-Homer1c, tdTomato-Gephyrin, HaloTag-Syb2, and mTagBFP2. **B,** puncta density of Homer1c:Syb2 and Gephyrin:Syb2 pairs at indicated timepoints. Two-way ANOVA (DIV, 0.66; puncta type, 0.0027; interaction, 0.74) with Fisher’s LSD post hoc. Mean ± SEM, N=3 cultures. **C,** excitatory/inhibitory ratios calculated from same cultures. Welch’s t-test (P=0.62). Mean ± SEM, N=3 cultures. **D,** representative traces for mEPSC/mIPSC ratio measurements recorded from DIV6, 10 and 16 primary hippocampal neurons. **E & F,** cell capacitance (E) and membrane resistance (F) from recordings at indicated timepoints. **G,** cumulative probability plot of mEPSC amplitude measurements. **H,** cumulative probability plot of mEPSC inter-event intervals. **I & J,** similar to G and H except for mIPSC measurements. **K,** average mEPSC/mIPSC amplitudes at indicated timepoints. **L,** average ratio of mEPSC/mIPSC amplitudes. One-way ANOVA with Tukey post hoc (ns – not significant). **M & N,** similar to **K** and **L** except for mEPSC/mIPSC frequency measurements. One-way ANOVA with Tukey post hoc (ns – not significant).

To examine functional maturation of developing excitatory and inhibitory synapses, we recorded spontaneous mEPSCs and mIPSCs during synaptogenesis (**Fig. 8D-N**). We monitored mEPSCs and mIPSCs from the same neurons by adjusting the holding potential (mEPSCs vH = -70 mV; mIPSCs vH = 0 mV) and analyzed their frequency and amplitude ratios at different maturation stages of the culture (DIV6, 10 and 16) (**Fig. 8D-N**). Cellular capacitance increased and membrane resistance decreased over time, consistent with previous reports and supporting maturation over culture time (**Fig. 8E & F**). While mEPSC amplitude modestly increased over culture time (**Fig. 8G**), mEPSC frequency increased and reached peak interevent interval by approximately DIV10 (**Fig. 8H**). mIPSC amplitude was stable between DIV6 and DIV10 and increased as neurons matured by DIV16 (**Fig. 8I**). mIPSC frequency progressively increased over time (**Fig. 8J**). We then calculated the ratios of mEPSC/mIPSC amplitude and frequency ratios (**Fig. 8K-N**) and observed no statistically significant differences between average mEPSC/mIPSC ratio measurements during culture maturation, suggesting that the development of excitatory and inhibitory spontaneous release is balanced as neurons mature.

## Discussion

Using live imaging approaches, we observed the dynamics between pre- and postsynaptic compartments in primary neuron cultures over extended time periods. These results indicate that subsets of synapses are spatially and temporally dynamic and continue to form and turnover even in relatively mature hippocampal neurons. Despite this turnover, excitatory and inhibitory synapses remain at an equilibrium over the time periods we examined. This balance is also preserved at the functional level, where the overall E/I ratios of spontaneous miniature release are maintained during maturation in cultured neurons. We postulate that intrinsic mechanisms possibly synchronize excitatory and inhibitory synapse assembly during circuit development. Two potential models may explain this synchronization. Temporal synchrony of intrinsic genetic programs may independently drive excitatory and inhibitory synapse maturation in parallel. Alternatively, regulation of excitatory and inhibitory synapse ratios may occur concurrently whereby signaling crosstalk between excitatory and inhibitory synapses dynamically balances excitation/inhibition and supports the functional outputs of neural circuits. These hypothetical mechanisms may help balance excitation/inhibition and support the functional outputs of neural circuits. Previous studies have demonstrated the role of synaptic plasticity and homeostatic plasticity on maintaining excitation/inhibition balance, demonstrating the intrinsic capacity for stimulus-dependent dynamic regulation^29,30^. Moreover, the activity state of a neuron/synapse including plasticity-inducing stimuli might alter the dynamic properties we observed and shift the balance of E/I synapse density. By tracking both excitatory and inhibitory synaptic structures in developing circuits, our approaches can be used in future studies to test these models.

Our live imaging studies enabled us to observe the motion and lifetime of synaptic compartments at key developmental timepoints. In addition to co-clustered pre- and postsynaptic compartments we observed abundant puncta without a nearby corresponding pre-or postsynaptic compartment, recapitulating what has been extensively observed in immunostaining of endogenous synaptic proteins. There are two possibilities for the molecular identity of these unpaired synaptic compartments. First, the absence of a cognate pre- or postsynaptic marker may be indicative of a synapse lacking the specific tagged proteins. For example, certain synaptic proteins may accumulate at specific synapses^31^, such that a given synaptic marker lacks the capacity to label all excitatory or inhibitory synapses. A second possible identity of unpaired pre- or postsynaptic proteins could be synapse precursor elements, which are subsequently stabilized upon recruitment to mature synaptic junctions. Indeed, we found that unpaired puncta exhibited distinct spatial and temporal characteristics, including shorter durations and faster speeds. This suggests recruitment at synapses could physically stabilize these protein clusters, prolonging their observable lifetime and physically restraining their motion. Available pre- or postsynaptic sites could also exist waiting for a viable partner but are eliminated without additional input, reducing average lifetime. Puncta lifetimes were also marker-specific, as seen with longer puncta duration of tagged Homer1c as compared to Syb2, indicating pre- and postsynaptic compartments lifetimes could be partially independent. Further studies are necessary to understand the full relationship between observed spatial dynamics and the functional consequences on developing circuits.

Our approaches also display several inherent limitations which will require future advances to overcome. While we focused on canonical and widely used pre- and postsynaptic markers as proxies for synaptic compartments, our tools do not assess the adhesive interaction between pre- and postsynaptic compartments. Combining our approaches with other systems relying on adhesion between pre- and postsynaptic cell adhesion molecules, such as SynView, GRASP, or SynapseShot^32–34^, will be important towards understanding further details of the dynamics of synapse assembly. Our approaches complement those based on trans-synaptic adhesion, considering the latter systems only exhibit signal after cell adhesion molecules contact, while our reporters can visualize synaptic components prior to forming a synapse. More generally, our observations support that some of the functional components of synapses are present in nascent pre- or postsynaptic compartments prior to their contact, including within growth cones. The live imaging tags in our tools can also be easily exchanged for a given experimental purpose or to introduce more robust labels as they are developed. Future studies combining our approaches with trans-synaptic adhesion reporters or genetically encoded activity indicators including GEVIs^35,36^ will help unravel the temporal sequences of synapse assembly and maturation.

Our live imaging was conducted in neuronal culture given its broad applicability and tractability for molecular studies. While this system provides a powerful and widely used reduced model for mechanistic questions, future advances will be necessary to employ these tools in combination *in vivo*. Our proof-of-concept studies support the utility of these tools *ex vivo*, establishing a foundation for future optimization of their use in neural circuits. Considering our approaches relied on reporter expression or CRISPR/Cas9 genome modifications, both of which require time for protein buildup to detectable levels, we were unable to visualize the very early stages of postnatal synaptogenesis. Synapses form in excess during postnatal development *in vivo*, followed by refinement and elimination of connections^37^. Future advances will be required to assess the events at each stage of postnatal synaptogenesis.

While tagged reporters for Homer1, Syb2 and Gephyrin are routinely used, our approaches provide several new features and applications compared to existing live imaging approaches, including intrabodies^38,39^. The lentiviral reporters and image tracking approaches are optimized for simultaneous monitoring of pre- and postsynaptic compartments, while currently reliable intrabody labeling approaches (FingRs) are available for postsynaptic proteins^40–43^. Furthermore, our viral approaches are designed to simultaneously image both excitatory and inhibitory synapses in the same neurons. While our TKIT studies focused on endogenous Gephyrin, the TKIT system can tag multiple different proteins in the same neuron. This raises the possibility of introducing distinct live imaging tags into different synaptic proteins in the same cell and monitoring them simultaneously. Ultimately, a combination of systems including viral tagging and intrabody labeling will be optimal for comprehensive evaluation of synapse dynamics during synaptogenesis.

Our work introduces new tools and methods that enable visualization of pre- and postsynaptic dynamics at high spatiotemporal resolution. We leveraged these new approaches to observe the spatial and temporal dynamics of individual synapses in neuronal cultures. Despite their continual temporal dynamics and turnover, excitatory and inhibitory synapse density remained balanced, which was further reflected by functional balance of excitatory and inhibitory spontaneous synaptic transmission. Future work using these tools and others will continue to illuminate the molecular mechanisms used by neural circuits to maintain steady state levels of synapses, and how these structures are dynamically generated and removed.

## Methods

### RESOURCE TABLE

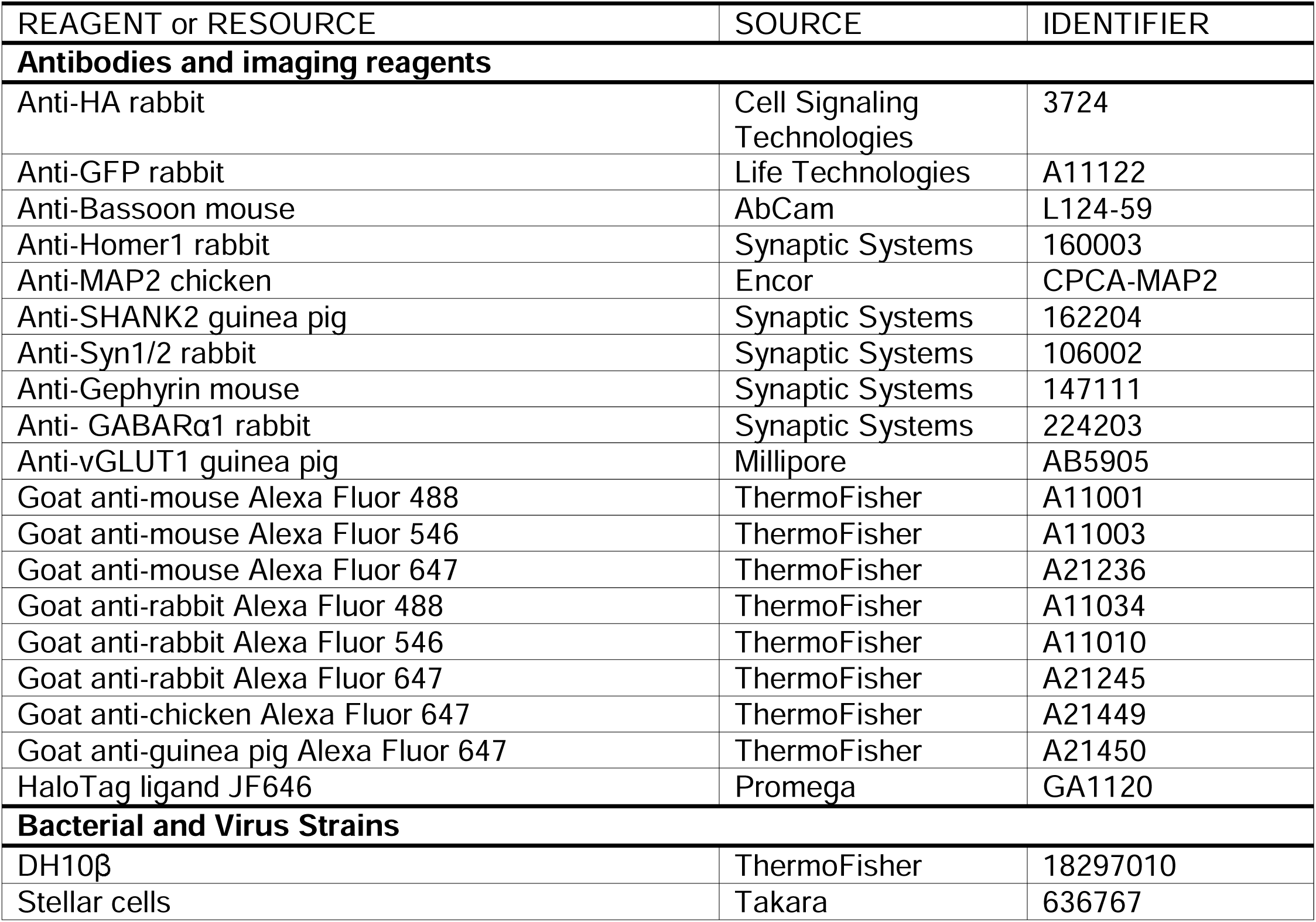
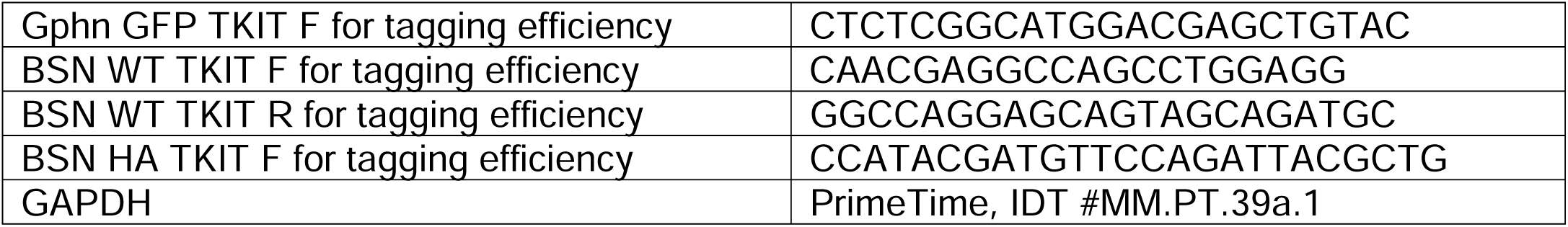
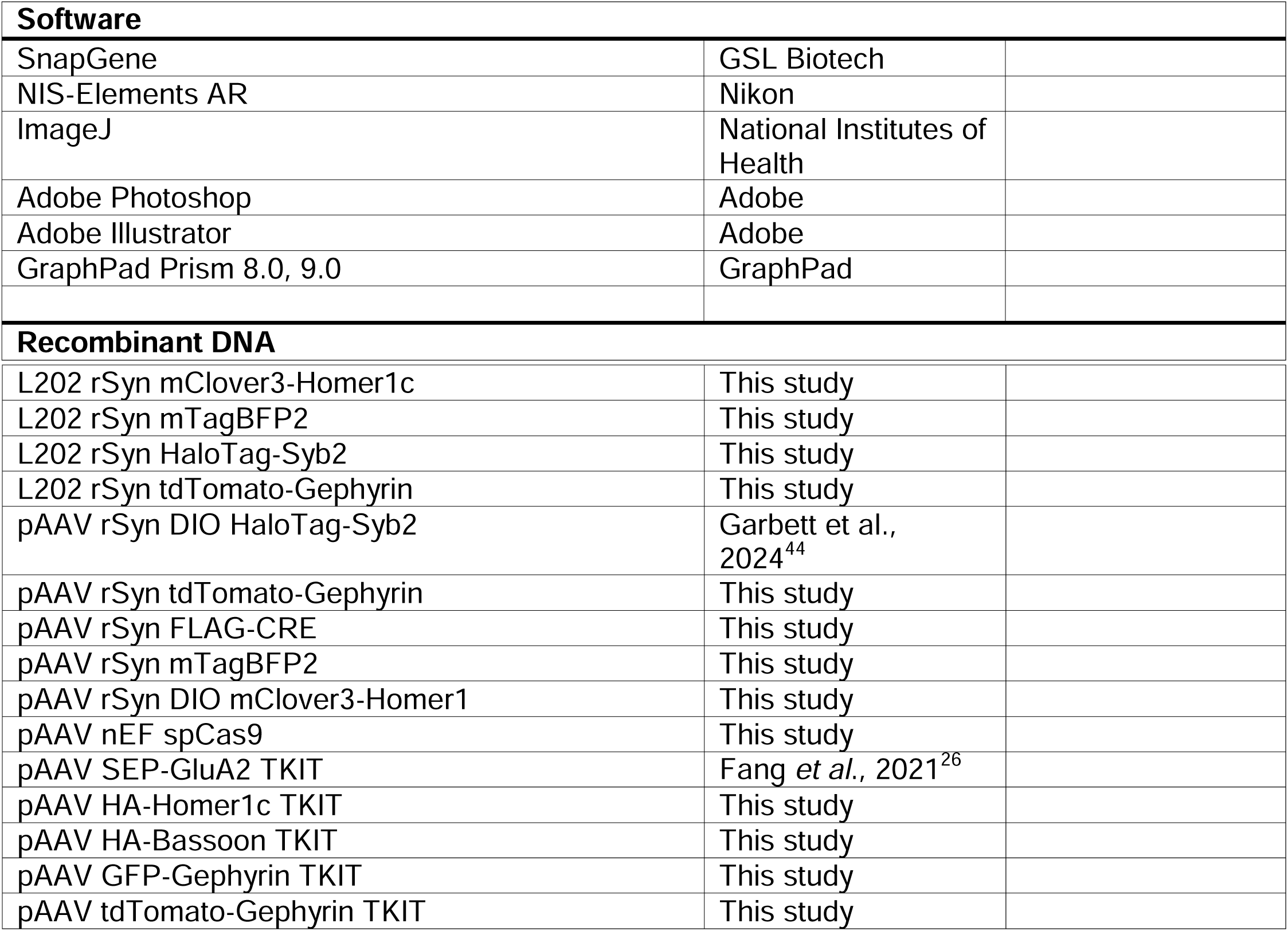

### RESOURCE AVAILABILITY

#### Lead contact

Further information and requests for resources and reagents should be directed to and will be fulfilled by the lead contact, Richard C. Sando.

#### Materials availability

All materials generated in this study will be openly shared upon request free of charge.

### EXPERIMENTAL MODELS AND SUBJECT DETAILS

#### Mice

C57BL/6J (Jax #000664) mice were used in this study. Mice were housed in groups of 2 to 5 on a 12 hr light/dark cycle with food and water *ad libidum* at the Vanderbilt Animal Housing Facility managed by the Division of Animal Care. All procedures conformed to National Institutes of Health Guidelines for the Care and Use of Laboratory Mice and were approved by the Vanderbilt University Administrative Panel on Laboratory Animal Care. Primary hippocampal cultures were generated from P0 pups.

#### Cell Lines

HEK293T cells (ATCC # CRL-11268) were maintained in DMEM (Gibco Cat# 11995065) containing 10% FBS (Gibco Cat# 16000044), 1X Penicillin-Streptomycin (Corning Cat# MT30002Cl) at 37°C and 5% CO_2_ for a maximum of 25 passage numbers.

#### Primary hippocampal cultures

For immunocytochemistry, hippocampal neurons were plated on a PDL-coated glass coverslip (#0, 12 mm, Carolina Biological Supply Company #633009) in 24-well plates (Genesee, cat # 25-107MP). PDL coating was accomplished using 50 mg/ml Poly-D-lysine (Gibco, Cat # A38904-01) coated for 1 hr to overnight at 37°C, followed by three washes with sterile dH_2_O and drying for 15 mins at RT. Mouse hippocampi were dissected from newborn mice and neurons were dissociated by papain (Worthington Biochemical Corporation, cat # LS003126) digestion for 20 min at 37° C, filtered through a 70 µm cell strainer (Corning, cat # 431751), and plated at density of 80,000 cells per dish/well. Plating media contained 5% fetal bovine serum (Life Technologies, cat # 16000044), B27 (Gibco, cat # 17504044), 1:50), 0.4% glucose, and 2 mM glutamine in 1x MEM. Culture media was exchanged 24 hr later (at DIV1) to growth medium, which contained 5% fetal bovine serum, B27, 2 mM glutamine in Neurobasal A (Gibco, cat # 10888022). Cytosine β-D-arabinofuranoside (Sigma, cat# C6645) was added to a final concentration of 2 µM on DIV3 in a 50% growth media exchange. Primary hippocampal cultures were infected with respective lentiviral/AAV conditions at DIV1.

#### Plasmids

Live imaging lentiviral reporters were encoded in a 3^rd^ generation lentiviral shuttle vector driven by the rat Synapsin promoter. The mClover3-Homer1c fusion contained mClover3 fused to the N-terminus of *Rattus norvegicus* Homer1 (NP_113895.1) separated by a glycine-serine linker (GGSGGGSGG). HaloTag-Syb2 was composed of HaloTag7 fused to the N-terminus of *Mus musculus* Syb2 (NP_033523.1) separated by a glycine-serine linker sequence (GSGGGG). The tdTomato-Gephyrin fusion contained tdTomato fused to the N-terminus of *Rattus norvegicus* Gephyrin (NP_074056.2) with a glycine-serine linker (GGSGGGSGG). TKIT plasmids were encoded in an AAV2 backbone harboring the tagged donor sequence together with a dual U6 promoter cassette delivering both sgRNAs. The donor sequence was designed as “flip and switch” as in Fang *et al*., 2021^26^. Each sgRNA was driven by a U6 promoter and followed by a gRNA scaffold sequence. All molecular cloning was conducted with the In-Fusion Assembly system (Takara #638948).

#### CRISPR Tag Knockin Design

Viral-based CRISPR tag knockins were designed according to the TKIT (Targeted KI with Two Guides) strategy in Fang *et al*., 2021^26^. The SEP-GluA2 TKIT vector was a kind gift from Drs. Richard Huganir and Alexei Bygrave. The SEP-GluA2 TKIT donor was subcloned into an AAV2 vector, and the same AAV2 vector was used for Homer1c, Gephyrin, and Bassoon KI constructs. sgRNAs were designed using the CRISPR sgRNA design tool from Integrated DNA Technologies. See Table 1 for all sgRNA sequences. For HA-Homer1c tagging, sgRNAs were designed to *Mus musculus* Ensembl variant 203 with a focus on the 5’ UTR and intron 1-2. For the Homer1c donor fragment, the HA tags were placed immediately following the start ATG. For HA-Bassoon tagging, the HA tag was inserted onto the N-terminus of Bassoon using a donor harboring HA immediately after the start ATG. For GFP-Gephyrin or tdTomato-Gephyrin tagging, the donor fragment was designed to insert the tag on the N-terminus of Gephyrin. CRISPR tagging vectors were cloned into an AAV2 backbone containing dual U6 promoters driving sgRNA1 and sgRNA2 for each target along with the donor tagging insert in a ‘flip-and-switch’ orientation as in Fang *et al*., 2021^26^, flanked by sgRNA sites. AAV encoding untagged spCas9 driven by the nEF promoter was used for all CRISPR modification experiments.

**Table 1.**
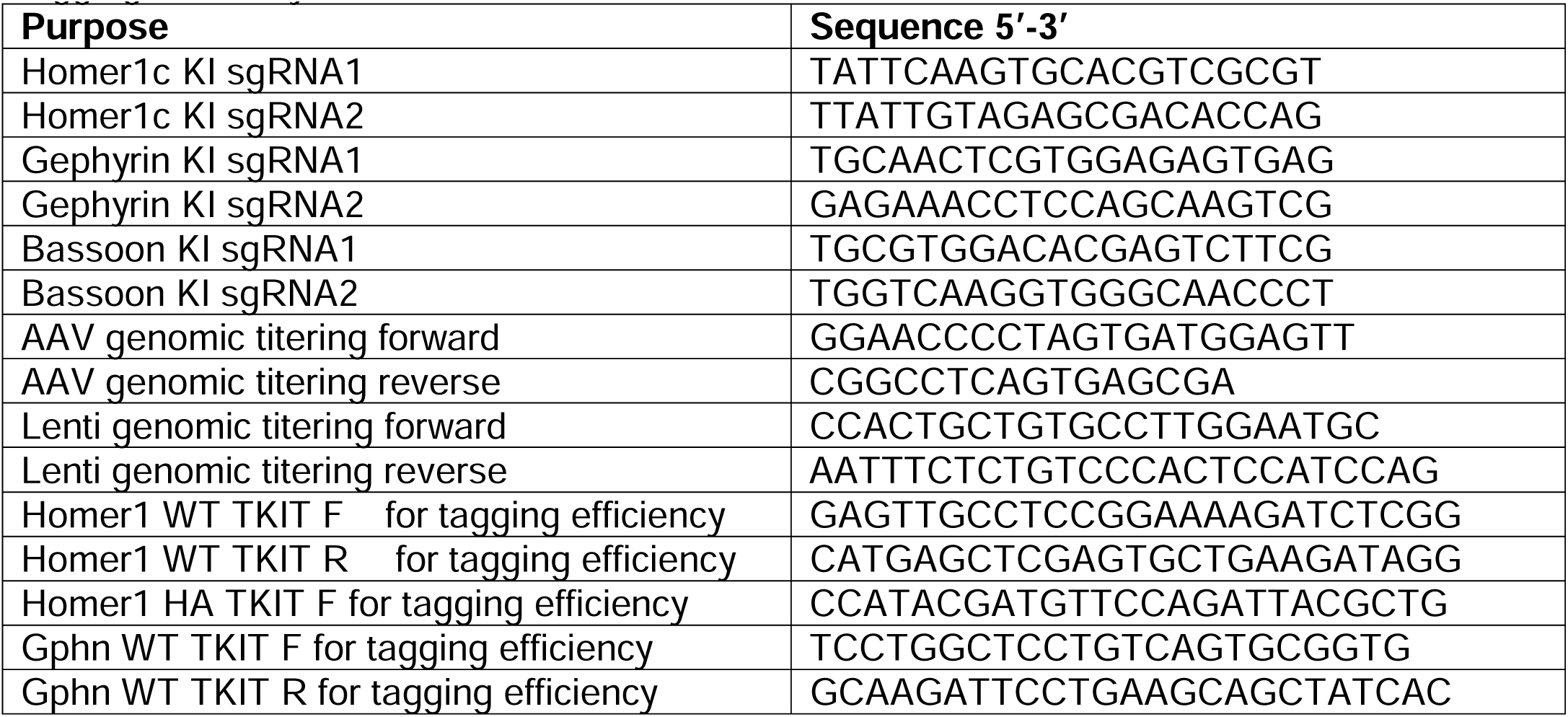
CRISPR KI sequences, AAV and Lenti titering primers, and primers for estimating the tagging efficiency of the TKIT method.

#### Antibodies

The following antibodies and reagents were used at the indicated concentrations: anti-HA rabbit (Cell Signaling Technologies, cat# 3724, 1:1,000), anti-HA mouse (Covance Cat# MMS101R; 1:1,000), anti-GFP rabbit (Life Technologies, cat# A11122, 1:1,000), anti-Bassoon mouse (AbCam, cat # L124-59, 1:1000), anti-Homer1 rabbit (Synaptic Systems, cat# 160 003, 1:5,000), anti-MAP2 chicken (EnCor Biotechnology, cat# CPCA-MAP2, 1:5,000), anti-SHANK2 guinea pig (Synaptic Systems, cat# 162204, 1:2,000), anti-Syn1/2 rabbit (Synaptic Systems, cat# 106 002, 1:5,000), anti-GABARα1 rabbit (Synaptic Systems, cat# 224203, 1:1,000), anti-Gephyrin mouse (Synaptic Systems cat# 147111, 1:2,000), anti-vGLUT1 (Millipore cat# AB5905, 1:1,000), corresponding fluorescently-conjugated goat secondary antibodies from Life Technologies (1:1,000).

#### Live imaging

For live imaging, hippocampal neurons were cultured in 35 mm culture dishes (MatTek Corporation, cat # P35G1.514C) with PDL-coated 1.5 mm glass bottom as described above at a density of 70,000 cells/dish in 1 mL of Growth Media. Prior to imaging, 0.2 µM JF646 Janelia Fluor® HaloTagTag® Ligand (Promega #GA1120) in 300 µL of the conditioned growth medium was applied to the cells for 2 hrs, followed by returning the cells back to the remaining 700 µL conditioned media. Images were acquired using a Nikon A1-R Eclipse Ti confocal microscope in resonant scanning mode with a 60x objective (Nikon #MRD01605, CFI60 Plan Apochromat Lambda, N.A. 1.4) and perfect focus operated by NIS-Elements AR acquisition software. Laser intensities and acquisition settings were established for individual channels using optimal LUT settings. Live images were acquired using multipoint acquisition to collect several cells/dish in a 1.5 µm Z-stack every 5 min at 4X averaging during a 15-hr period (181 time frames total per cell) while normal growth conditions were maintained (37°C, 5% CO_2_, 90% humidity) using a Tokai Hit stage incubator (Tokai Hit STXG Incubation System).

#### Live Image Analysis

Timeseries images with three z-steps of 0.5 µm step size were flattened with the Extended depth of focus algorithm in NIS-Elements. Crops of 92.07 x 92.07 microns (400x400 pixels) were made representative areas of original images, and background was subtracted using a custom FIJI/ImageJ Python script. Timeseries images were corrected with a custom-built python drift correction package. Live imaging analysis and synaptic puncta tracking was performed with a custom-built python package. For each condition, 1-5 areas containing multiple neurites were analyzed from 3-5 independent cultures.

##### Drift Correction

To isolate local biological motion from global microscope drift, we employed a tiled phase-correlation approach designed to remain robust to local motion. Image channels were combined into a composite image and divided into a 10 x 10 tile grid. Tile emptiness was estimated using Otsu threshold mask coverage, and tiles with less than 1% mask coverage were excluded from drift estimation. The vector of motion between corresponding tiles between frame reference pairs was estimated with phase correlation. For each frame reference pair, a consensus cluster of coordinated x- and y-shift vectors was identified with DBSCAN clustering (ε=2 px), and the mean within this consensus cluster was used as the drift estimate. Drift was estimated relative to 10 distributed reference frames, and the final drift trace for each frame was computed as the equally weighted mean of the reference-specific drift estimates after re-basing each estimate to the first frame according to the following equation:

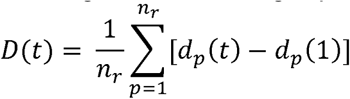

where *D(t)* is the final 2D drift vector at time *t*, *d_p_(t)* is the drift vector estimated at time *(t)* relative to reference frame *p*, and *n_r_* is the number of reference frames. These drift estimates were then used to generate stabilized images for downstream analysis.

##### Puncta Segmentation

To determine puncta positions and centroids, we segmented puncta using a custom python package. Briefly, Laplacian of Gaussian blob detection was used to identify diffraction-limited spots of typical synaptic puncta sizes (0.52-1.44 µm diameter). These spots were used as a region-of-interest for a local seed growing method with high and low thresholds derived from local background statistics. Touching segmented puncta were merged but re-evaluated by watershed based peak re-detection when multiple local maxima were present. Final puncta positions were determined using an intensity-weighted centroid within puncta segmentation masks. To ensure temporal consistency of detections, a second detection sweep was run with relaxed threshold in regions where a punctum was predicted in an adjacent frame but not initially detected. Puncta were assigned intensity-based quality scores using a Gaussian mixture model, and low-confidence detections were removed.

##### Pairing Analysis

To enforce one-to-one assignment of pre- and postsynaptic puncta, we modeled synapse pairing with a linear assignment problem (LAP), which had been previously used for synaptic colocalization^45^. Pairing costs were computed from a hybrid distance and segmentation-overlap term. The cost function between puncta *i* and *j* was computed as the following:

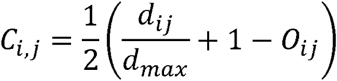

Where *d_ij_* is the distance between puncta *i* and *j*. *d_max_* was the maximum allowed pairing distance. O*ij* is the segmentation area of overlap score between puncta *i* and *j*, or the intersection over union of the segmented masks. Candidate pairs were only considered if *d_ij_* < *d_max_*, where *d_max_* = 2 µm. Final synapse positions were defined as the midpoint of the paired pre-and postsynaptic puncta positions.

##### Puncta density calculations

To determine puncta density, dendrites containing both pre- and postsynaptic components were manually traced in ImageJ/FIJI. Traced lines were smoothed to prevent overestimation of dendrite length due to slight local deviations. Puncta were segmented and paired as previously, and puncta counts along the line within 30 pixels (3.45 µm) were counted. Individual puncta and paired puncta densities are reported as puncta / µm dendrite. Given paired puncta types A paired with B, the paired puncta percentage was calculated as

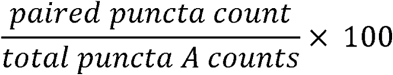

while total pairing counts of punctum types A and B were

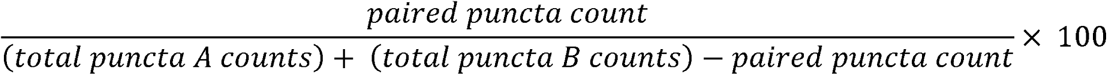

In total, 1-4 dendrites per area with 1-5 areas per replicate culture, totaling 20-24 dendrites per condition. Puncta densities were averaged per culture for statistical analysis.

##### Puncta Tracking Analysis

Synaptic puncta tracking and paired puncta tracking was performed using a global min-cost flow (MCF) solver, which has been previously used for synapse tracking^46^. Tracking was performed essentially as previously described for MCF multi-object tracking^47^. Candidate trajectories were constructed using four edge types: birth edges from a source node to each detection, transit edges through each detection, link edges between detections in subsequent frames, and death edges from detections to a sink node. A zero-cost bypass edge from source to sink allowed flow to pass unused, such that tracks were only created when favored for the lowest cost solution.

Candidate links were generated between detections in consecutive frames or across a single missed frame. For a temporal gap of Δ*t* frames, candidate links were restricted by a maximum displacement gate of *d_max_* √Δ*t*, where *dmax* = 1 μm, and each detection was connected to at most 10 nearest-neighbor candidates per gap. For each candidate link between detections *i* and *j*, the link cost was defined as

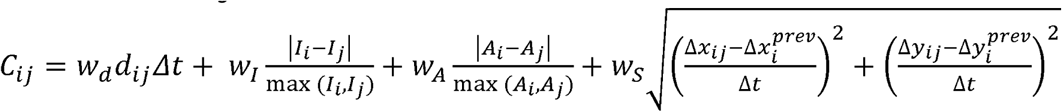

Where *d_ij_* is centroid displacement in micrometers, *I* is the puncta intensity, *wI* is the intensity weight, *A* is the puncta area, *wA* is the area weight, and the final term is the magnitude of the change in velocity of the puncta from the previous frame pair to current pair *ij*, with a motion smoothness weight *wS*. The corresponding weights were *w_d_* = 4.0, *wI* = 0.1, *wA* = 0.1, and *wS* = 1.0. Thus, links with larger displacement, larger changes in intensity or area, or larger changes in velocity were penalized. Each detection transit cost was defined as:

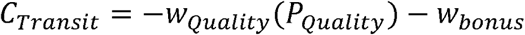

Where P*_Quality_* is a detection quality score (0-1) derived from fitted intensity distributions. W*_Quality_* = 10 and W*_Bonus_* = 0.1. Birth and death costs were each set to 15, and the total cost of a track was the sum of its birth, transit, link, and death costs. The MCF algorithm (Google OR-Tools) determined the globally optimal flow configuration for each movie and each object class (pre-, postsynaptic and pairs) to generate final tracks.

##### Puncta track categorization

Because tracking allowed single frame gaps, we analyzed the total number of observed detection frames in a track divided by the total track span in frames and referred to this as the detection persistence. For all analyzed tracks, we only considered tracks where detection persistence >75%. Tracks were categorized as Stable (>95% duration, 790 min), Stable-like (95%-20% duration, 160-790 min), and Transient (< 20% duration, 160 min). Stable-like tracks were further categorized by Eliminated (present in first 10 frames, absent in last 10 frames), Nascent (absent in first 10 frames, present in last 10 frames) and Turnover (remaining Stable-like, effectively absent in first 10, and absent in last 10). For pre- and postsynaptic tracks, the paired fraction was determined for each track, which was the total count of paired puncta per track divided by the track length. Puncta tracks were defined as paired when paired fraction > 0.75, and unpaired when paired fraction < 0.05.

#### RT-qPCR

Total RNA was isolated from cultured neurons at DIV14 using RNAqueous™-Micro kit (Invitrogen, AM1931), and then used for cDNA synthesis with Super Script IV (Invitrogen, cat # 18090050) with random hexamers. qPCR with PowerUp SYBR Green Master Mix (Applied Biosystems, cat # A25742) was used for quantification of tagged transcripts and total (tagged + non-tagged) transcripts, and the housekeeping gene GAPDH. The quantification was done by generating a standard curve with plasmids harboring the donor DNA to calculate copy number. The primer pairs are listed in Table 2.

**Table 2.**
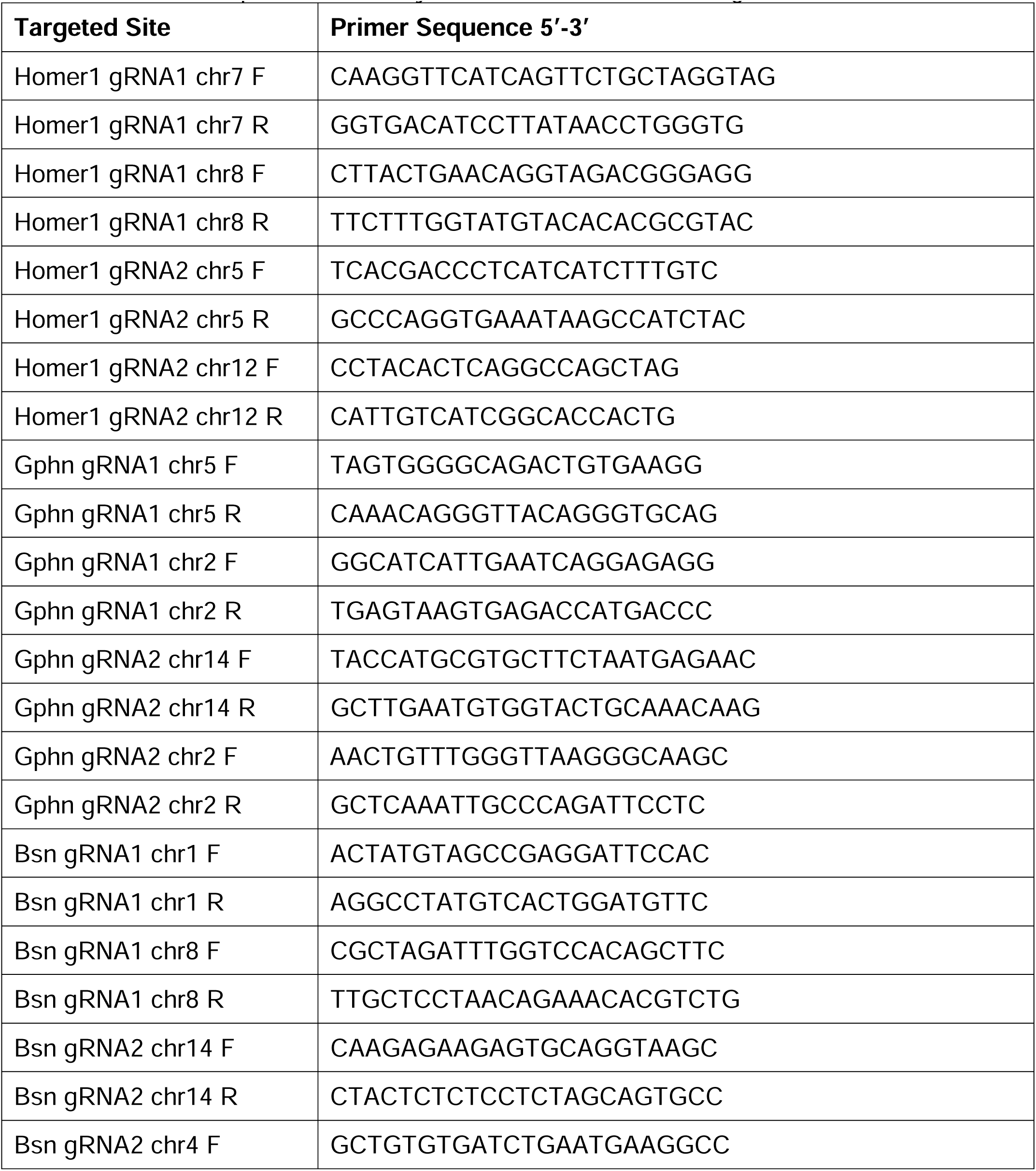
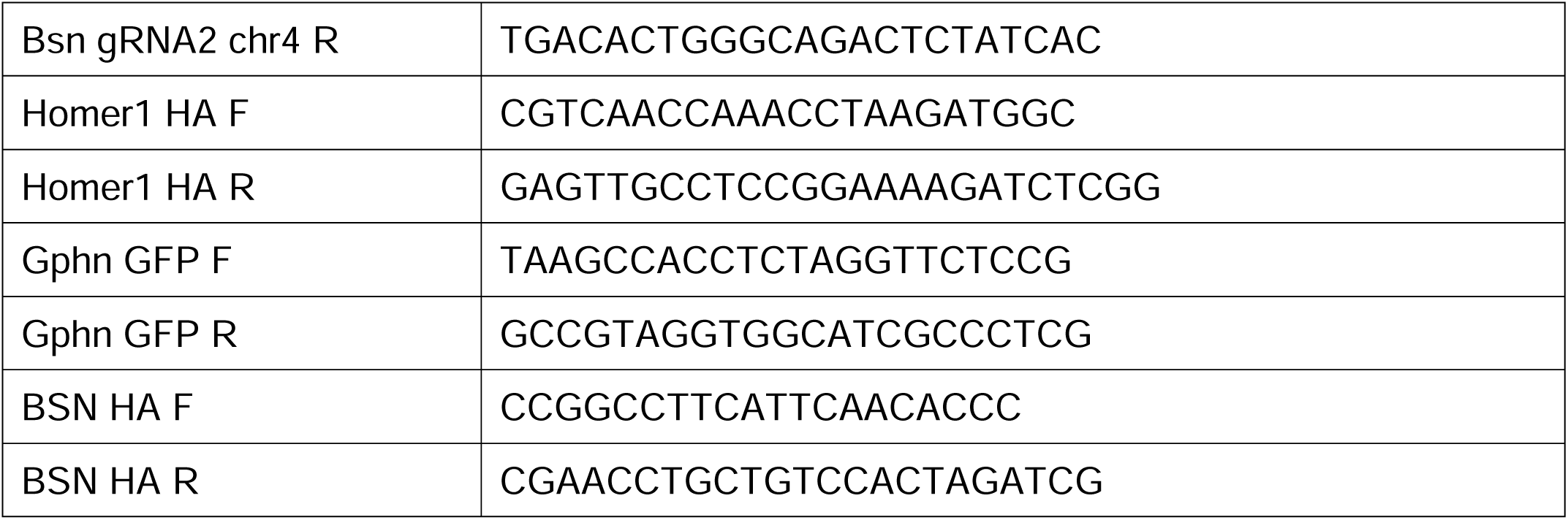
Primers sequences for analysis of CRISPR/CAS9 off-target modifications.

#### Analysis of CRISPR/CAS9 off-target effects

Site specific sequencing was used to screen for off-target CRISPR effects. The top two off-target sites for each gRNA used were identified with the CRISPR-Cas9 guide RNA design checker tool by Integrated DNA Technologies. Primers were designed to amplify the selected sites from genomic DNA and the resulting amplicons were Sanger sequenced (Eurofins Genomics). Genomic DNA was isolated from primary neuronal cultures with a tag introduced at a single gene using the Qiagen DNeasy Blood and Tissue Kit (Qiagen #69504). PCR reactions were conducted with DreamTaq (Thermo Scientific #K1081) and 10 ng genomic DNA template. The primer sequences are listed in Table 2 and the sequence results from the potential off-target sites surrounded by 10 nucleotides are presented in Fig. S6 along with all necessary details pertaining to the genomic location.

#### Immunocytochemistry

Cover glass (#0, 12 mm, Carolina Biological Supply Company #633009) was placed into 24-well plates and coated for 2 hrs with 100 µL of 50 µg/mL poly-D-lysine (Gibco #A38904-01) in the 37°C tissue culture incubator. Excess poly-D-lysine was removed, coverslips were washed 3x with sterile ddH_2_O and dried for 30 mins. Neurons were plated at a density of 70,000 cells/well. At DIV10-14, cells were washed briefly once with PBS, fixed with 4% PFA (Electron Microscopy Science Cat# 15714)/4% sucrose/PBS for 20 min at 4°C, and washed 3 x 5 mins in PBS. Samples were permeabilized in 0.2% Triton X-100/PBS for 5 mins at room temperature and then transferred to blocking buffer (4% BSA (Sigma Cat# 10735086001)/3% normal goat serum (Jackson Immunoresearch #005000121)/PBS). Samples were incubated in blocking buffer for 1 hr, and subsequently incubated with diluted primary antibody in blocking buffer for 2 hrs at room temperature. Samples were then washed 5 x 5 mins in PBS, incubated with fluorescently conjugated secondary antibodies diluted in blocking buffer for 1 hr at room temperature, washed 5 x 5 mins in PBS and mounted on UltraClear microscope slides (Denville Scientific Cat# M1021) using 10 µL ProLong Gold antifade reagent (Invitrogen, #P36930) per coverslip. Samples were dried at RT in the dark prior to imaging.

#### Immunohistochemistry and HaloTag labeling

Mice were briefly anesthetized with isoflurane and transcardially perfused with 10 mL room temperature heparinized (10 U/mL, Sigma #H3393) oxygenated ACSF, followed by 25 mL room temperature 4% PFA/PBS. The brains were post-fixed 2 hrs at 4°C, washed with PBS, cryoprotected in gradients of 10% sucrose/PBS, 20% sucrose/PBS, and 30% sucrose/PBS, rapidly embedded in OCT (Fisher # 23730571), and sliced on a cryostat at 25 µm. For immunohistochemistry, the 25 µm thick, free-floating sections were washed 1 x 5 mins with PBS/0.1% Triton X-100, blocked for 1 hr at room temperature in blocking buffer (4% BSA (Sigma Cat# 10735086001)/3% normal goat serum (Jackson Immunoresearch #005000121)/0.1% Triton-X100/PBS) and incubated overnight at 4°C with primary antibodies diluted in blocking solution. Primary antibodies included anti-GFP rabbit (1:500, Life Technologies #A11122). Primary antibody incubation was followed by 3 x 5 min washes in PBS/0.1% Triton X-100 and 2 hr room temperature incubation with corresponding fluorescently labeled secondary antibody (goat anti-rabbit AlexaFluor488, ThermoFisher #11034) in blocking buffer. Samples were labeled with DAPI (Sigma Cat#10236276001) diluted into PBS/0.1% Triton X-100 for 15 mins, washed 5 x 5 mins with PBS/0.1% Triton X-100, and mounted on glass slides coated in 0.1% Triton-X100/PBS, dried briefly, and covered with ProLong Gold antifade reagent (Invitrogen, #P36930) and coverglass (Corning Cat#2980-246). For HaloTag-Syb2 labeling in fixed slices, injected animals were perfused as above, post-fixed in ice-cold 4%PFA/PBS for 2 hrs, cryoprotected in gradients of 10% sucrose/PBS, 20% sucrose/PBS, and 30% sucrose/PBS, and subsequently rapidly frozen in OCT (Fisher # 23730571). Coronal sections at 25 µm thickness were obtained via cryostat, washed 1x with PBS, and incubated overnight at 4°C with 200 nM HALO ligand JF646 (Promega #GA1120). Sections were subsequently counterstained with DAPI/PBS for 15 mins at room temperature, washed 4 x 5 mins with PBS, and mounted onto UltraClear microscope slides (Denville Scientific Cat# M1021) using ProLong Gold antifade reagent (Invitrogen, #P36930).

#### Lentivirus production for culture experiments

Lentiviruses were packaged in HEK293T cells from ATCC (CRL-11268). For lentiviral production, co-transfection of the expression shuttle vector and the three helper plasmids (pRSV-REV, pMDLg/pRRE and vesicular stomatitis virus G protein (VSVG)) was done with FuGENE6 (Promega E2691) using 2.5 µg of each plasmid per 9.6 cm^2^. Lentiviral-containing medium was collected 48 hr after transfection, briefly spun down 5,000 xg for 5 mins for removal of cellular debris and then stored at 4°C. The LV genomic titer was estimated using PowerUp™ SYBR™ Green Master Mix for qPCR (Applied Biosystems, A25742) with the following primers: F - ccactgctgtgccttggaatgc, and R - aatttctctgtcccactccatccag. Shuttle plasmids at 10x serial dilutions (1x10^5^ - 1x10^9^ copies/mL) were used for generating a standard curve. After quantification, the LVs were directly applied to primary neuron culture medium.

#### Adeno-associated virus production

For production of AAVs, five 10-cm^2^ plates of HEK293T cells at 90% confluency were transfected via the calcium phosphate method. For transfections, 100 µg of each plasmid (pHelper, pDJ serotype, and AAV shuttle plasmid) was mixed to a volume of 6.75 mL dH_2_O, and 0.75 mL of 2.5 M CaCl_2_ was added. The DNA/CaCl_2_ mixture was added dropwise to 7.5 mL 2× HBS, pH 7.05 (274 mM NaCl, 10 mM KCl, 1.5 mM Na_2_HPO_4_, 7 H_2_O [dibasic], 12 mM dextrose, and 42 mM HEPES) while vortexing gently. The DNA/CaPO_4_ mixture was incubated at room temperature for 20 min, and then 3 mL was added dropwise to each plate. Cells were washed 1× with prewarmed PBS 24 hr after transfection, and medium was replaced with fresh complete DMEM (DMEM + 10% FBS + 1X Pen/Strep.). Cells were harvested 72 hrs after transfection by 1× wash with PBS followed by addition of dissociation buffer (PBS/10 mM EDTA). A cell scraper was used to facilitate detachment, and cell suspensions were subsequently centrifuged at 1,500×g for 15 min at 4°C. Cell pellets were resuspended in 4 mL freezing buffer (150 mM NaCl, 20 mM Tris, pH 8.0, and 2 mM MgCl_2_), snap-frozen in 70% ethanol/dry ice for 15 min, and rapidly thawed at 37°C. After three subsequent rounds of freeze/thaws, the cell suspension was incubated in 50 U/mL Benzonase nuclease (Sigma; Cat# E1014) for 30 min at 37°C. Samples were subsequently centrifuged at 3,000×g for 30 min. Supernatant was applied to the surface of an iodixanol gradient (15%, 25%, 40%, and 60%) and ultracentrifuged at 80,000×g for 2 hr at 4°C in Seton Scientific Polyclear thin walled centrifuge tubes (7030). The 40% iodixanol gradient was harvested by puncturing the side of the tube with a sterile needle attached to a 10 mL syringe, added to 10 mL PBS/1 mM MgCl_2_, and concentrated in centricon concentrating tubes (100,000 MWCO; Millipore; UFC0910024), which were equilibrated with PBS/MgCl_2_. After 3 subsequent washes with 1x DMEM (Gibco Cat# 11995065), samples were concentrated to 100 µL, aliquoted, and stored at −80°C. The AAV particle number was estimated using PowerUp™ SYBR™ Green Master Mix for qPCR (Applied Biosystems, A25742) with primers targeting the ITR: F - ggaacccctagtgatggagtt, and R – cggcctcagtgagcga (AddGene). Packaged shuttle plasmid 10x serial dilutions (1x10^5^ - 1x10^9^ copies/mL) were used for generating a standard curve.

#### Confocal Imaging of fixed samples

Images were acquired using a Nikon A1r resonant scanning Eclipse Ti2 HD25 confocal microscope with a 10x (Nikon #MRD00105, CFI60 Plan Apochromat Lambda, N.A. 0.45), 20x (Nikon #MRD00205, CFI60 Plan Apochromat Lambda, N.A. 0.75), and 60x (Nikon #MRD01605, CFI60 Plan Apochromat Lambda, N.A. 1.4) objectives, operated by NIS-Elements AR v4.5 acquisition software. Laser intensities and acquisition settings were established for individual channels and applied to entire experiments. Image analysis was conducted using Nikon Elements and ImageJ. Brightness was adjusted uniformly across all pixels for a given experiment for Figure visualization purposes. Image channels were pseudocolored for Figure visualization purposes.

#### Stereotactic Injections

Stereotactic injections were performed on P21-25 mice anesthetized with 1-5% vaporized isoflurane. Analgesia was provided with a preoperative subcutaneous injection of 0.05 mg/g meloxicam (PCAA 55-4476), followed by additional doses at 24 and 48 hours post-procedure. Mouse heads were shaved and cleaned with Betadine followed by 70% ethanol. Heads were secured to a stereotactic rig (Stoelting Digital Lab Standard with mouse and neonate adaptor) and lubricant was applied to eyes (Puralube Vet Ointment). A small incision was made through the scalp with sterilized tools. All viral solutions were injected using a beveled glass pipette (Warner Instruments G120F-4) connected to a Hamilton 1701RNR 10 µL Syringe (#80065) with 18-gauge syringe needle (Hamilton RN NDL 18-gauge S #7804-06), completely backfilled with mineral oil. A syringe pump (World Precision Instruments #SP100I) delivered 0.2 µL per injection at 0.9 µL/hr. The following viral combinations were injected into the CA1 region of the hippocampus: AAV hSyn tdTomato-Gephyrin + AAV DIO HaloTag-Syb2; AAV hSyn DIO mClover3-Homer1c + AAV hSyn FLAG-Cre; and AAV GFP-Gephyrin TKIT + AAV nEF Cas9 + AAV Syn mTagBFP2. Coordinates used for bilateral CA1 injections were A/P - 1.80 mm, M/L +/- 1.20 mm, and D/V -1.35 mm. The following viral combination was injected into the CA3 region of the hippocampus: AAV hSyn DIO HaloTag-Syb2 + AAV hSyn FLAG-Cre. Coordinates used for bilateral CA3 injections were A/P -1.80 mm, M/L +/- 2.30 mm, D/V -2.10 mm. To prevent capillary action, the injection pipette was left at the injection site for 10 mins post-injection, then slowly raised +0.05 mm dorsally and remained for an additional 2 mins before being slowly removed. Incisions were sutured and sealed with Vetbond tissue adhesive (3M #1469SB) and mice were removed from rig. Efficiency and localization of viral injections was confirmed by a Nikon Eclipse FN1 microscope.

#### Electrophysiology

Primary hippocampal cultures were made as described above. Whole cell voltage patch clamp was performed on cells on DIV6, DIV10, and DIV16. Patch pipettes were pulled from borosilicate glass capillary tubes (World Precision Instruments; Cat# TW150-4) using a PC-10 pipette puller (Narishige). Pipette resistance filled with intracellular solution ranged between 3- 4 MOhm. Intracellular solution consisted of (in Mm) 125 Cesium gluconate, 5 CsCl, 8 NaCl, 8 HEPES, 1 EGTA, 2 MgATP, 0.3 Na_2_GTP, 2 Phosphocreatine. Final pH was adjusted to 7.25-7.3 with 1M HCl and 300 Osm. External solution contained (in mM) 140 NaCl, 5 KCl, 2 CaCl_2_, 0.8 MgCl_2_, 10 HEPES, and 10 glucose (pH 7.4, adjusted with NaOH). 1 µM tetrodotoxin was added to external solution to isolate miniature spontaneous synaptic activity. Synaptic currents were monitored with a Multiclamp 700B amplifier (Molecular Devices) synchronized with Clampex 11.2 data acquisition software. Electrophysiological data were digitized with Digidata 1550B (Molecular Devices). Cells were patched and recorded at -70 mV to isolate miniature excitatory currents for 2.5 minutes. After recording, the holding potential was slowly increased in 5 mV steps to 0 mV to isolate miniature inhibitory currents. Cells were allowed to stabilize for 1-2 minutes before recording inhibitory currents for 5 minutes. Recording parameters were assessed before recording excitatory currents, before recording inhibitory currents, and after inhibitory recordings to ensure the cell remained stable and healthy throughout the recording. Synaptic currents were sampled at 10 kHz and analyzed using Clampfit 11.2 software. Miniature events were analyzed using the template matching search and a minimal threshold of 5 pA, with each event visually inspected for inclusion or rejection.

### QUANTIFICATION AND STATISTICAL ANALYSIS

#### Statistics

All data represent the results of at least three independent biological replicates, as indicated within each Figure Legend. Statistical significance was determined using the two-tailed Student’s t-test, one-way ANOVA with following *post hoc* Tukey tests for multiple comparisons, or two-way ANOVA with following *post hoc* Tukey tests for multiple comparisons, as indicated in the Figure Legends. Paired tests were used when assessing within culture differences in speed or duration for concurrently imaged puncta. Data analysis and statistics were performed with Microsoft Excel, GraphPad Prism 8.0 and GraphPad Prism 9.0.

## Acknowledgements

We thank Stefanie Wieckert (Nikon) for assistance with imaging and imaging analysis procedures. We thank Dr. Alexei Bygrave (Huganir laboratory, Johns Hopkins University) for advice regarding the TKIT tagging approach. We thank Dr. Ege Kavalali (Vanderbilt University) and all members of the Sando laboratory for critical feedback on the manuscript and study. This study was supported by grants from the NIH (R00MH117235 and DP2MH140134 to RS) and Alfred Sloan Foundation (Sloan Fellowship in Neuroscience to RS).

## Author Contributions

K. Garbett performed all imaging experiments. J. Allen created puncta tracking computational image analysis methods. K. Garbett and R. Sando performed all molecular cloning. K. Garbett, J. Allen, J. Lopez, C. Smith and R. Sando performed analysis and interpretation. J. Lopez performed electrophysiology. C. Smith conducted stereotactic injection studies. K. Garbett, J. Allen and R. Sando designed the study. K. Garbett, J. Allen, and R. Sando wrote the manuscript with input from all authors.

## SUPPLEMENTAL DATA FIGURES and FIGURE LEGENDS

**Figure S1:**
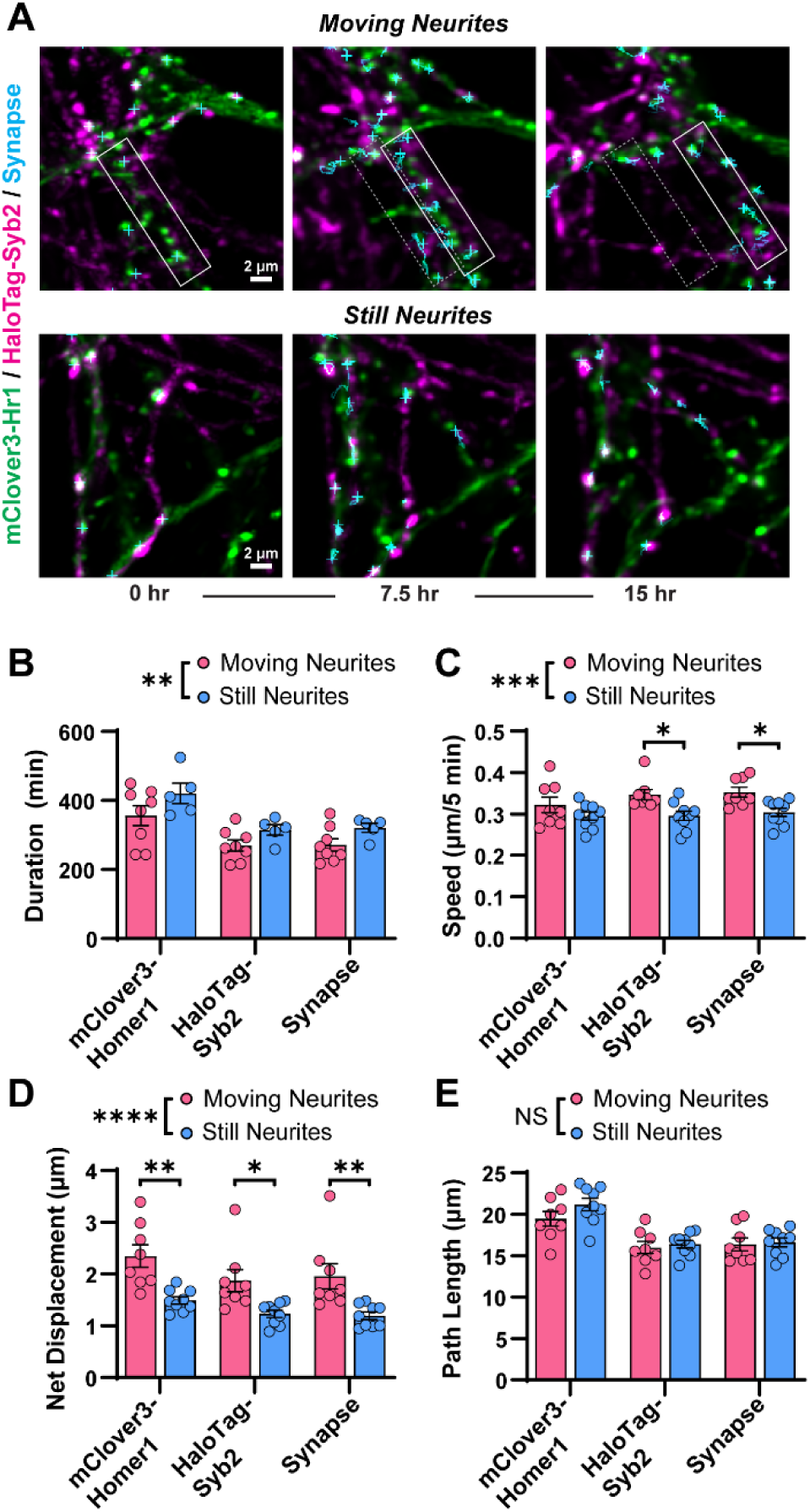
Tracking neuron cultures with heterogenous motion. **A**, representative images of cultures classified as having moving neurites (*top*) and still neurites (*bottom*). Gray and white boxes in moving neurites depict a neurite that moves throughout the imaging session. **B**, duration of puncta in movies binned into moving or still neurites for mClover3-Homer1c, HaloTag-Syb2, or synapse pairs. Two-way ANOVA (Moving-Still, P=0.0074; Puncta-type, P=0.0002; interaction P=0.9) with Tukey’s post hoc. **C,** as in B but with puncta speed. Two-way ANOVA (Moving-Still, P=0.0001; Puncta-type, P=0.28; interaction P=0.57) with Tukey’s post hoc. **D,** as in B but with net displacement (start to end distance). Two-way ANOVA (Moving-Still, P=<0.0001; Puncta-type, P=0.044; interaction P=0.79) with Tukey’s post hoc. **E,** as in B, but with path length (total distance traveled). Two-way ANOVA (Moving-Still, P=0.17; Puncta-type, P=<0.0001; interaction P=0.52) with Tukey’s post hoc.

**Figure S2:**
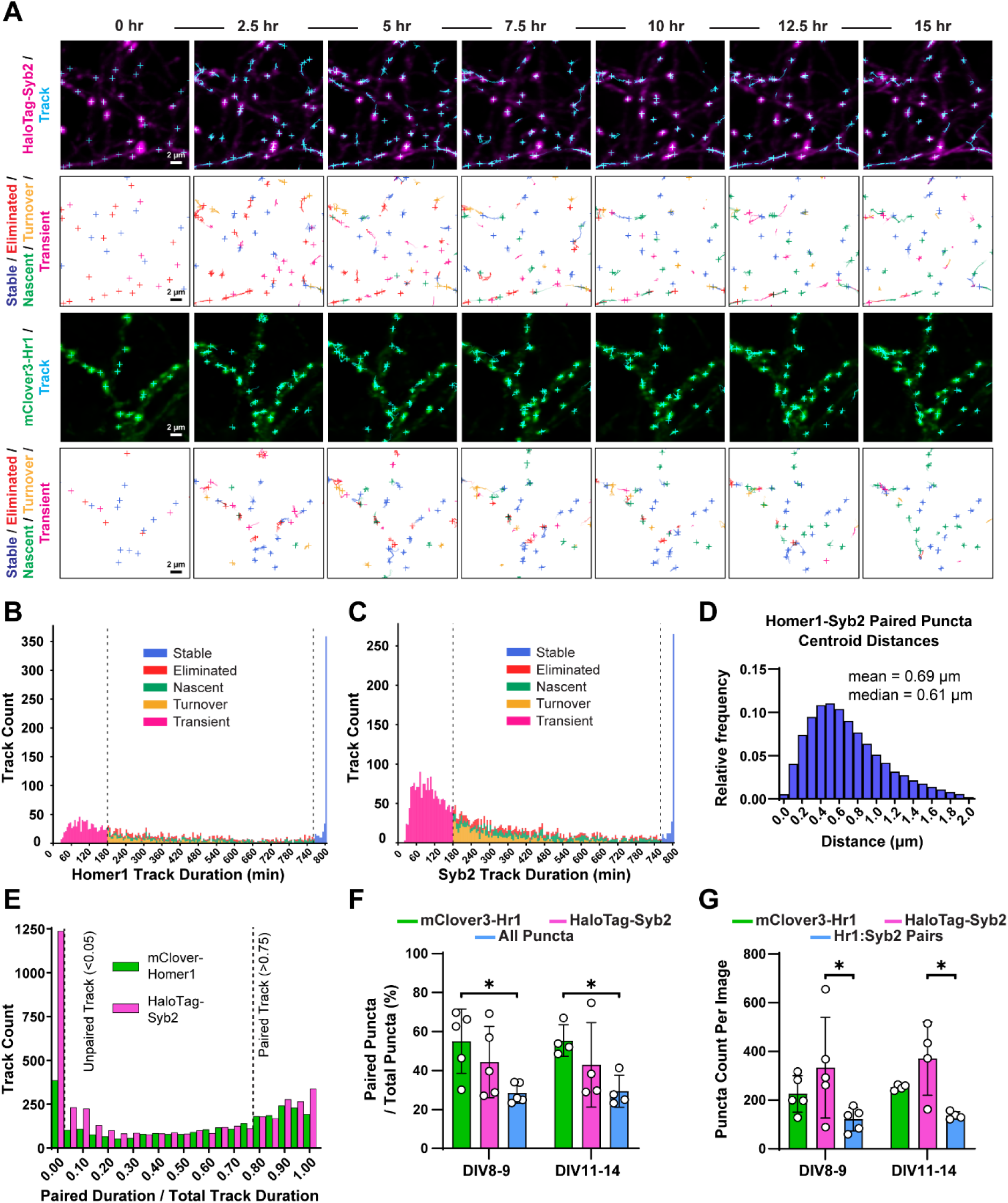
Additional parameters for excitatory synapse dynamics across development, related to Figures 2 and 3. **A**, example HaloTag-Syb2 (*top*) and mClover3-Homer1c (*bottom*) tracks with track categories (as in Figure 2). **B & C,** histogram of track duration for mClover3-Homer1c (B) and HaloTag-Syb2 (C). **D,** histogram of inter-puncta centroid distances for mClover3-Homer1c and HaloTag-Syb2 puncta. **E,** histogram of puncta pairing duration for mClover3-Homer1c (green) and HaloTag-Syb2 (magenta). Tracks were classified as unpaired (pairing duration < 5% total time) or paired (pairing duration > 75% total time). **F**, pairing analysis of data shown in Figure 3. Counts of paired puncta / the total puncta of that type are shown. For all puncta, this represents all puncta pairs relative to total number of puncta of any type. Two-way ANOVA (DIV, P=0.99; puncta type, P=0.0035; interaction, P=0.98) with Tukey’s post hoc. Mean ± SEM, N=4-5 independent cultures. **G**, total puncta count of all puncta types for mClover3-Homer1c, HaloTag-Syb2, and puncta pairs. Two-way ANOVA (DIV, P=0.55; puncta type, P=0.0019; interaction, P=0.98) with Tukey’s post hoc. Mean ± SEM, N=4-5 independent cultures.

**Figure S3:**
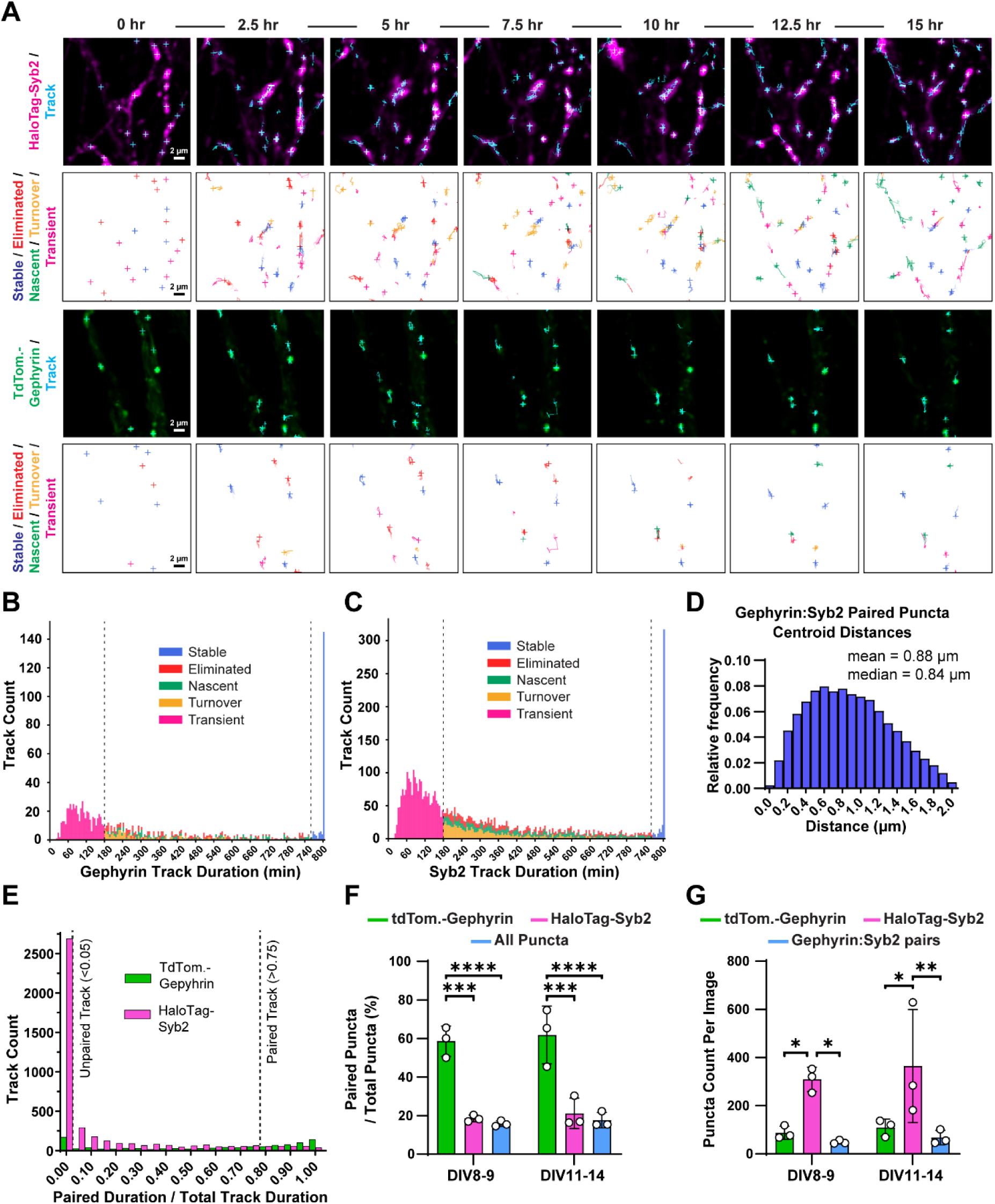
Additional parameters for inhibitory synapse dynamics across development, related to Figure 5. **A**, example HaloTag-Syb2 (*top*) and tdTomato-Gephyrin (*bottom*) tracks with track categories (as in Figure 2). **B & C,** histogram of track duration for tdTomato-Gephyrin (B) and HaloTag-Syb2 (C). **D,** histogram of inter-puncta centroid distances for tdTomato-Gephyrin and HaloTag-Syb2 puncta. **E,** histogram of puncta pairing duration for tdTomato-Gephyrin (green) and HaloTag-Syb2 (magenta). Tracks were classified as unpaired (pairing duration < 5% total time) or paired (pairing duration > 75% total time). **F,** pairing analysis of data shown in Figure 5. Counts of paired puncta / the total puncta of that type are shown. For all puncta, this represents all puncta pairs relative to total number of puncta of any type. Two-way ANOVA (DIV, P=0.50; puncta type, P<0.0001; interaction, P=0.98) with Tukey’s post hoc. Mean ± SEM, N=4-5 independent cultures. **G,** Total puncta count of all puncta types for tdTomato-Gephyrin, HaloTag-Syb2, and puncta pairs. Two-way ANOVA (DIV, P=0.51; puncta type, P=0.0008; interaction, P=0.94) with Tukey’s post hoc. Mean ± SEM, N=4-5 independent cultures.

**Figure S4:**
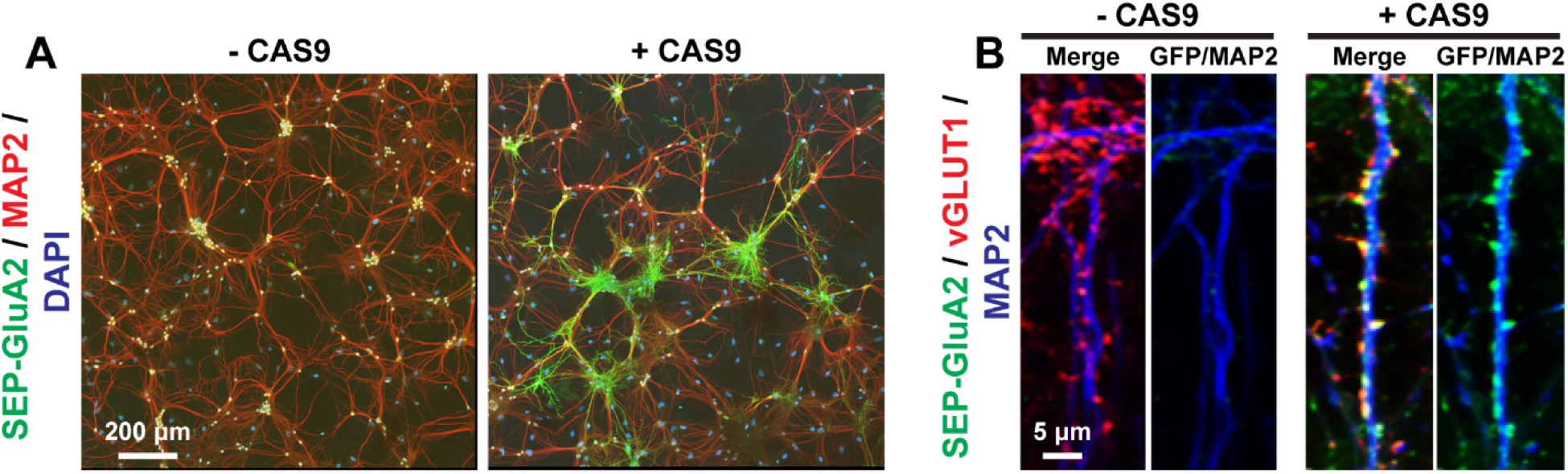
CRISPR/Cas9 tagging of endogenous GluA2 using the TKIT approach. **A & B**, example of the efficacy of the TKIT tagging approach with the SEP-GluA2 tagging constructs from Fang *et al*., 2021^22^. **A,** low magnification overviews illustrating the prevalence of SEP-tagged GluA2 neurons across a neuronal culture. **B**, high magnification representative dendritic stretches of endogenously tagged GluA2. Cells were co-labeled for GFP together with excitatory presynaptic vGLUT1 and the somatodendritic marker MAP2.

**Figure S5:**
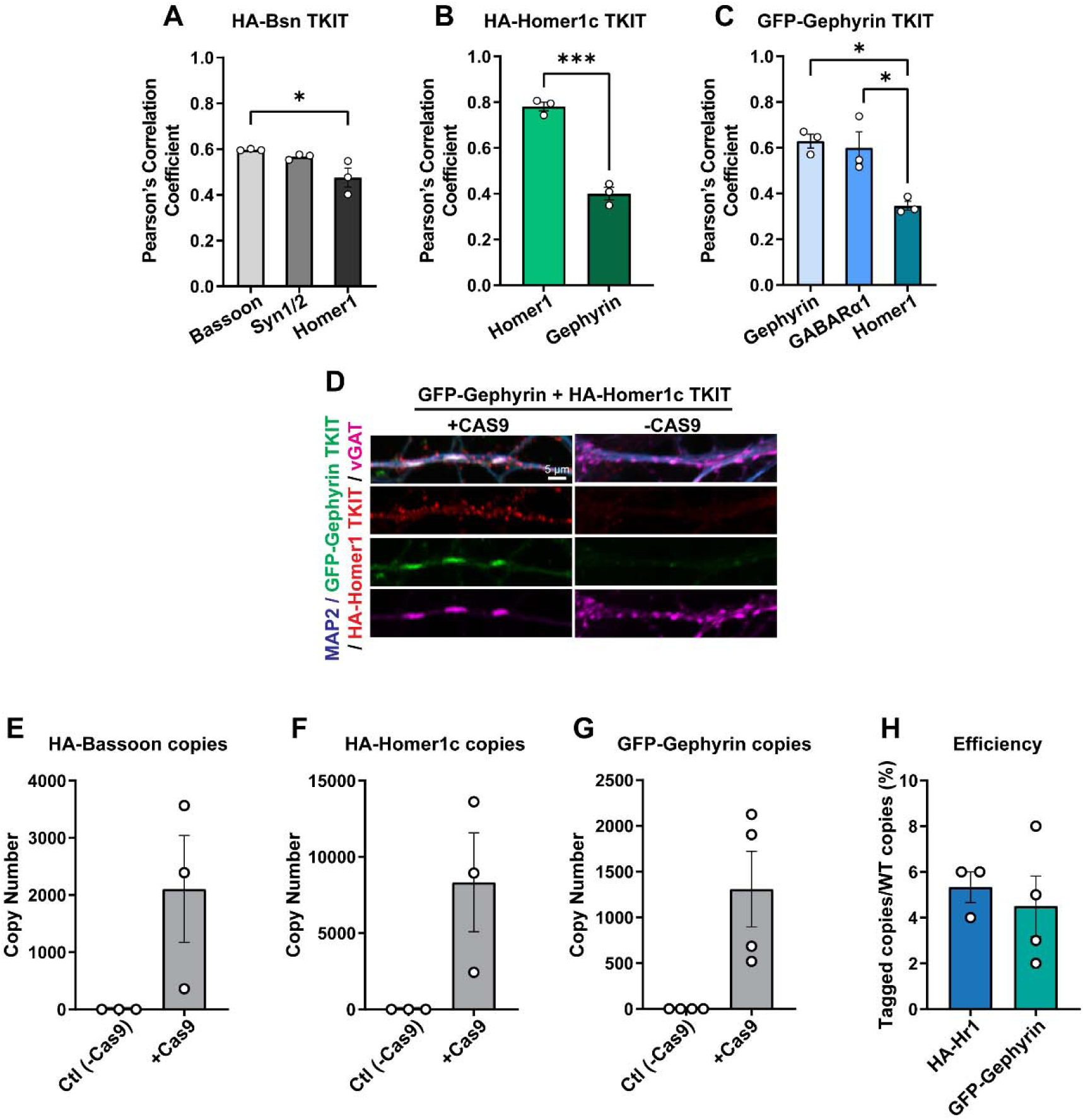
Additional characterization of CRISPR/Cas9 TKIT-mediated approach to label endogenous pre- and postsynaptic markers. **A-C**, quantification of Pearson’s correlation coefficient from results in Figure 6. **A,** Pearson’s correlation coefficient measurements of HA-tagged endogenous presynaptic Bassoon co-immunostained for indicated synaptic markers. **B,** similar to **A**, except for the HA-tagged Homer1c TKIT. **C,** similar to **A**, except for GFP-tagged Gephyrin. **D,** example TKIT tagging of both Gephyrin and Homer1c. **E-H,** RT-qPCR analysis of CRISPR/Cas9 tagging efficiency. **E,** qPCR probes detecting the HA-tagged Bassoon transcript are only amplified when neurons are co-transduced with AAVs encoding the DNA donor/sgRNAs, and Cas9. Copy numbers were calculated via standard curve qPCR using a plasmid containing the tagged sequence. **F,** similar to **E,** except for HA-tagged Homer1c transcript. **G,** similar to **E,** except for GFP-tagged Gephyrin transcript. **H,** efficiency estimates based on standard curve qPCR with primers detecting tagged Homer1c or Gephyrin transcript specifically, or primers detecting both tagged and non-tagged transcripts. Numerical data are means ± SEM from 3-4 independent culture replicates indicated as open circles. Statistical significance was assessed via two-tailed t-test or one-way ANOVA with post hoc Tukey tests (*, p<0.05; ***, p<0.001).

**Figure S6:**
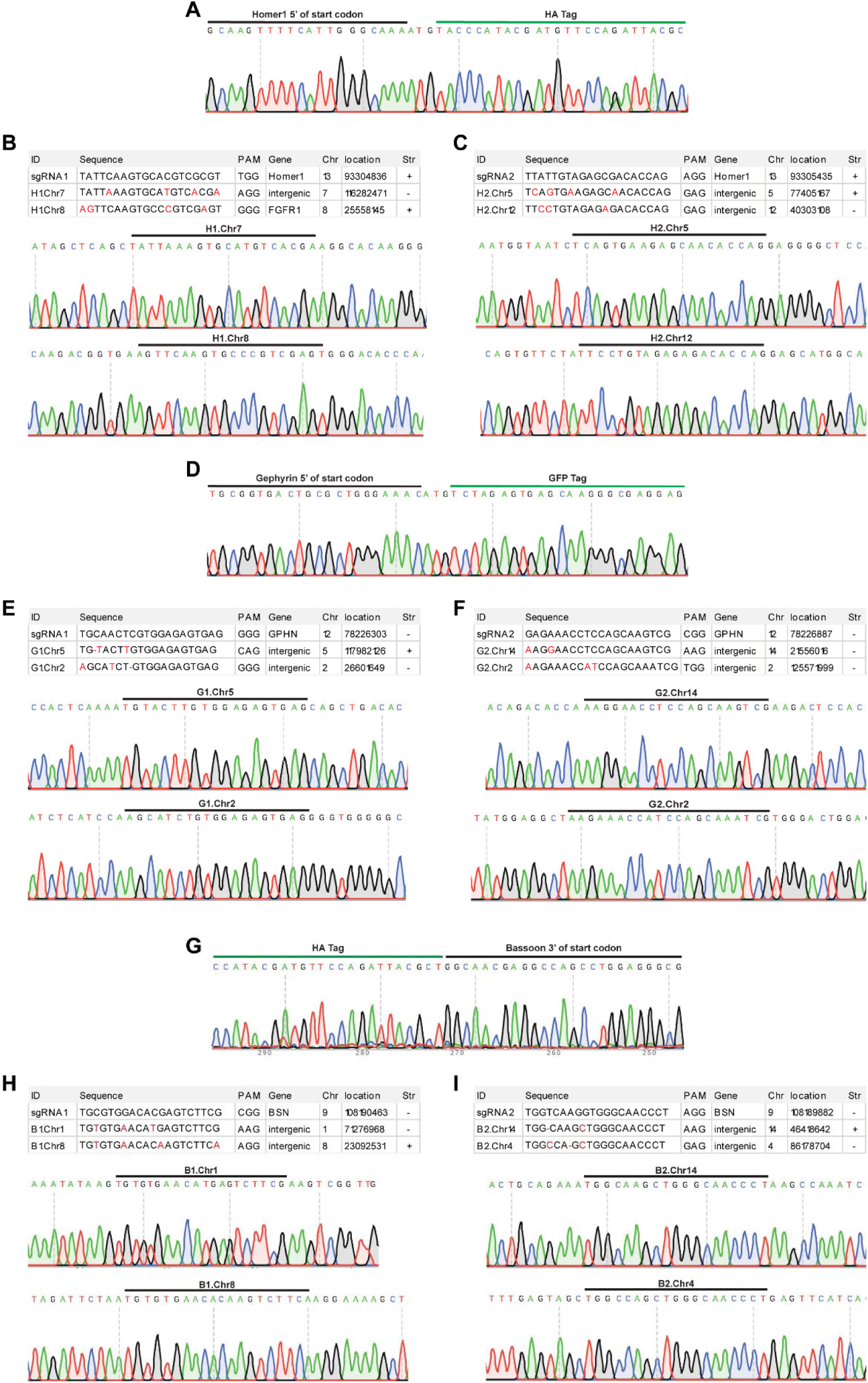
Analysis of TKIT off-target effects. **A**, chromatogram depicting HA insertion into the 5’ of endogenous Homer1c. **B,** Homer1c sgRNA1 sequence compared to the top two predicted off-target sites. Chromatograms depict sequencing of off-target sites. **C,** similar to **B,** except for Homer1c sgRNA2. **D,** chromatogram depicting GFP insertion into the 5’ of endogenous Gephyrin. **E,** Gephyrin sgRNA1 sequence compared to the top two predicted off-target sites. Chromatograms depict sequencing of off-target sites. **F,** similar to **E,** except for Gephyrin sgRNA2. **G,** chromatogram depicting HA insertion into the 5’ of endogenous Bassoon. **H,** Bassoon sgRNA1 sequence compared to the top two predicted off-target sites. Chromatograms depict sequencing of off-target sites. **I,** similar to **H,** except for Bassoon sgRNA2. Data depicts representative chromatograms from three independent experiments.

**Figure S7:**
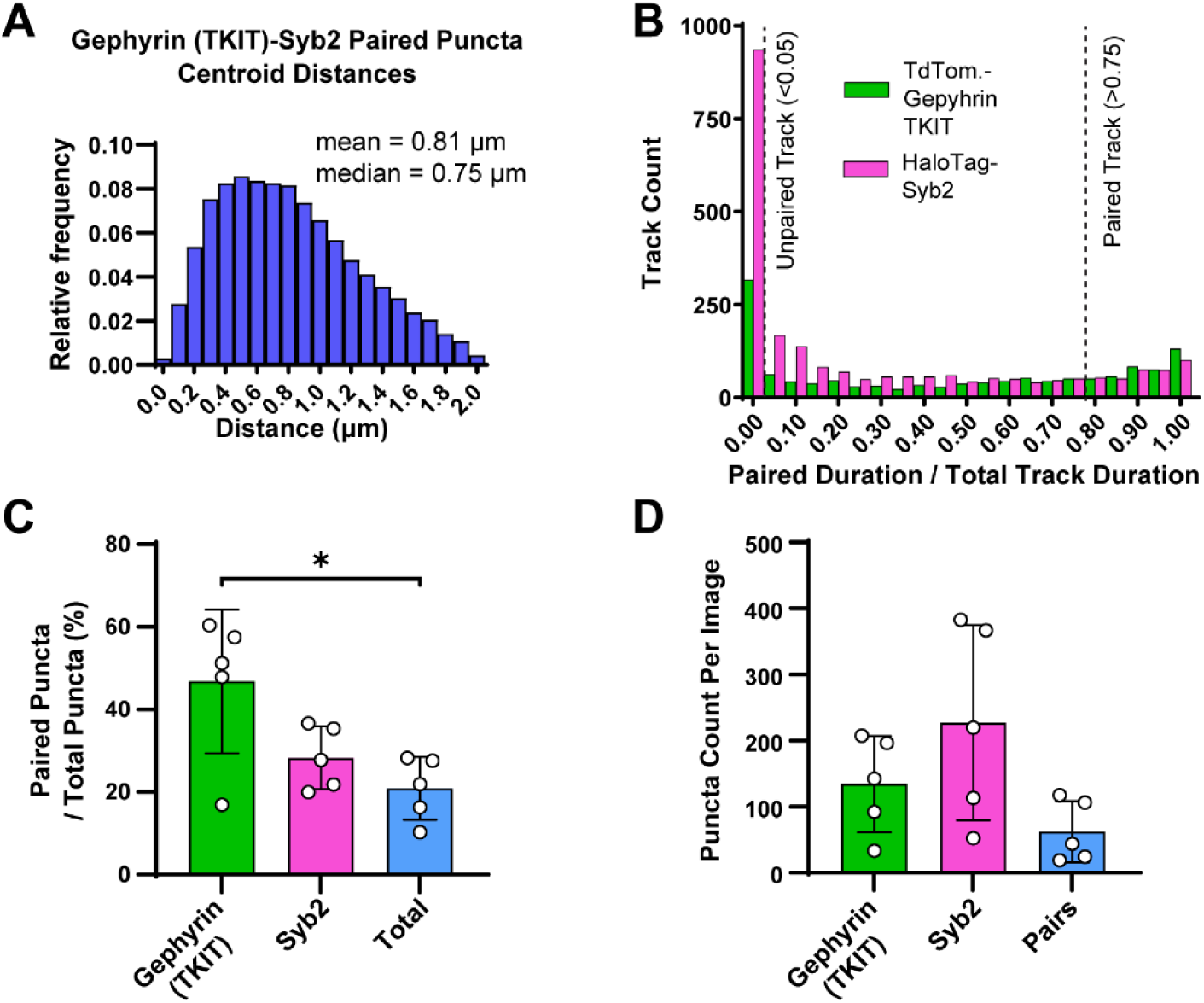
Additional parameters for tracking endogenous Gephyrin, related to Figure 7. **A**, histogram of inter-puncta centroid distances for endogenous tdTomato-Gephyrin and HaloTag-Syb2 puncta. **B,** histogram of puncta pairing duration for endogenous tdTomato-Gephyrin (green) and HaloTag-Syb2 (magenta). Tracks were classified as unpaired (pairing duration < 5% total time) or paired (pairing duration > 75% total time). **C,** pairing analysis of data shown in Figure 7. Counts of the number of paired puncta / the total puncta of that type are shown. For all puncta, this represents all puncta pairs relative to total number of puncta of any type. One-way ANOVA (P=0.013) with Tukey’s post hoc. Mean ± SEM, N=5 independent cultures. **D,** total puncta counts of all puncta for endogenous tdTomato-Gephyrin, HaloTag-Syb2, and puncta pairs. Kruskall-Wallis test (P=0.07). Mean ± SEM, N=5 independent cultures.

