## Supplementary material for "Stable excitatory-inhibitory synapse balance despite dynamic turnover": Movie Figure Legends

**Supplemental Movie 1: Representative live neuronal cultures expressing lentiviral mClover3-Homer1c, HaloTag-Syb2, and mTagBFP2.** Imaged at DIV9. Same culture as shown in Figure 1J.

**Supplemental Movie 2: Comparison of drift corrected to un-stabilized images.** First, drift corrected movie is shown, followed by uncorrected. Finally, side-by-side comparison of corrected and uncorrected on a region of the image. Graph above shows estimated per-frame drift (µm). Lateral jitter can be observed when estimated drift spikes.

**Supplemental Movie 3: Representative tracking of HaloTag-Syb2, mClover3-Homer1c, and synapse pairs with track categorization.** Crosshairs show current position while trails show previous positions of previous 30 frames. Individual channel tracks followed by track categorization are as described in Figure 2.

**Supplemental Movie 4: Representative live neuronal cultures expressing lentiviral tdTomato-Gephyrin, HaloTag-Syb2, and mTagBFP2.** Imaged at DIV12.

**Supplemental Movie 5: Representative live neuronal cultures expressing endogenous (TKIT CRISPR/Cas9-labeled) tdTomato-Gephyrin, lentiviral HaloTag-Syb2 and mTagBFP2.** Imaged at DIV13.
